# Ultrafast tyrosine-based cell membrane modification *via* diazonium salts: a new frontier for biomedical applications

**DOI:** 10.1101/2025.11.15.688583

**Authors:** Mohammed Bouzelha, Karine Pavageau, Sarah Renault, Enora Ferron, Nicolas Jaulin, Maia Marchand, Dimitri Alvarez-Dorta, Roxanne Peumery, Mathieu Scalabrini, Héloïse Delépée, Allwyn Pereira, Elise Enouf, Mathieu Cinier, Oumeya Adjali, Sébastien G. Gouin, Christelle Retière, Mickaël Guilbaud, David Deniaud, Mathieu Mével

**Affiliations:** Nantes Université, TaRGeT, Translational Research for Gene Therapies, CHU Nantes, INSERM, UMR 1089, 44000, Nantes, France; INSERM UMR1307, CNRS UMR6075, CRCI²NA, Team 12, Nantes, France; Etablissement Français du Sang, 44011, Nantes, France; Capacités, SAS, 44200, Nantes, France; Nantes Université, CNRS, CEISAM, UMR 6230, 44000, Nantes, France; Affilogic, SAS, 44300, Nantes, France

**Keywords:** Cell membrane, Tyrosine, Bioconjugation, Click-chemistry, Targeting, Cancer

## Abstract

In this study, we present an ultrafast, efficient, and broadly applicable strategy for cell membrane modification *via* tyrosine bioconjugation using diazonium salt derivatives. This chemical approach enables both one-step and two-step functionalization of adherent, suspension, and primary cells with diverse ligands, including imaging probes, carbohydrates, biotin, and proteins, without inducing cytotoxicity or immune activation.

Membrane engineering through bioconjugation is emerging as a valuable tool in biomedical research, given the cell membrane’s central role in signaling, transport, and cell-cell interactions. Compared to traditional glyco-engineering methods that often require multiday incubations and can cause cellular stress, our approach achieves precise and high-density grafting in less than one hour, with improved reproducibility and biological compatibility.

Importantly, we demonstrate that this strategy can be applied to a variety of cell types, encompassing both immortalized cell lines and primary cells, notably Peripheral Blood Mononuclear Cells (PBMCs) and human effector cells such as natural killer (NK) cells, to enhance their cytotoxic function against EGFR-positive cancer cells via surface-conjugated Nanofitin. Moreover, we show that the bioconjugated signal diminishes over time due to cell division, offering a self-limiting alternative to permanent genetic modifications such as CAR-T, thereby mitigating risks associated with overly prolonged immune activation.

Finally, the feasibility of storing pre-functionalized cells at -80 °C expands the practicality of this platform for future ready-to-use applications. Together, these features make our method a compelling alternative to current technologies for applications in targeted therapy, diagnostics, and cell-based immunotherapy.

## 1. Introduction

The modification of cell surfaces is a rapidly expanding field, driven by diverse approaches such as genetic engineering, metabolic glyco-engineering (MGE), and bioconjugation. Genetic engineering, particularly the introduction of peptides or proteins, is a widely utilized technique with profound implications for both research and therapeutic applications. A prominent example is CAR-T cell therapy,[1] where T cells are engineered to recognize and eliminate cancer cells, providing effective treatments for diseases like acute lymphoblastic leukemia.[2,3] These modifications are typically achieved using viral vectors,[4,5] which efficiently deliver genetic material encoding the chimeric antigen receptor (CAR), ensuring stable expression and precise targeting of cancer cells.[6] However, the use of viral vectors carries inherent risks, including unintended genomic integration that may lead to tumorigenesis or adverse immune responses.[7,8] In addition, the generation of CAR-T cells is a lengthy and costly process, generally requiring 2–4 weeks of *ex vivo* manipulation and specialized GMP-compliant facilities. These limitations underscore the need for safer and faster complementary strategies. In contrast, metabolic glycan engineering (MGE) offers a non-genetic strategy for cell surface functionalization by exploiting the cell’s endogenous glycan biosynthesis pathways (**Figure 1**).[9] This approach enables the incorporation of modified monosaccharides into membrane glycans, facilitating applications such as targeted labeling and cellular imaging. Since its inception in the 1980s, MGE has evolved to include unnatural sugar analogs,[10] bioorthogonal chemistries,[11] and an expanded range of applications.

**Figure 1:**
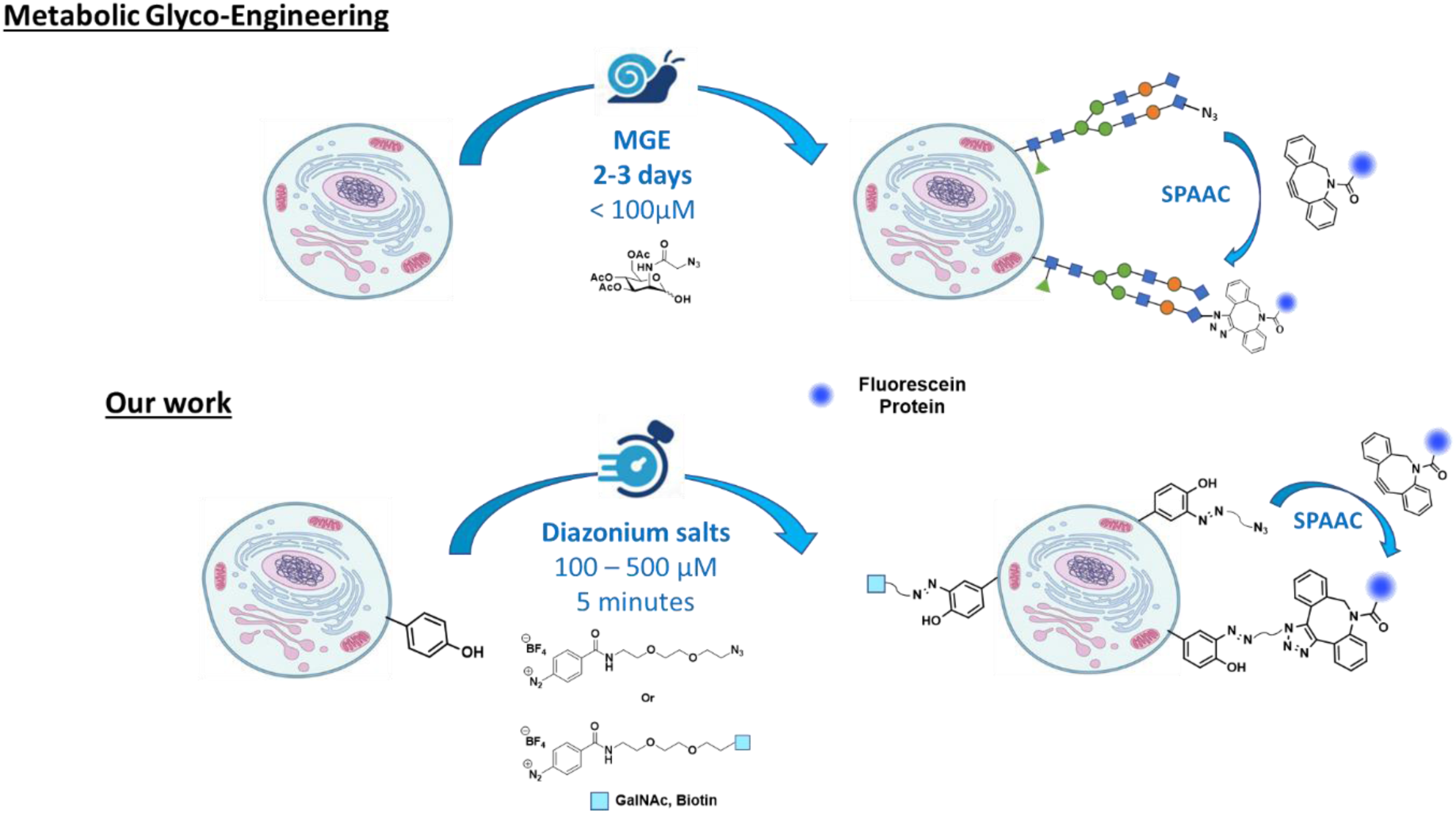
MGE and diazonium salts bioconjugation cell strategies. MGE: cells are incubated with Ac_4_ManNAz for 2–3 days, enabling azide incorporation into membrane glycans, followed by strain-promoted azide-alkyne cycloaddition (SPAAC) reaction with DBCO-PEG_4_-FAM for fluorescent labeling. Diazonium salt: cells are incubated for 5 min with diazonium salt ligands (N₃, GalNAc, or biotin), followed, in the case of N₃, by a SPAAC reaction with DBCO-PEG4-FAM or DBCO-Nanofitin for detection or functionalization.

However, despite its versatility, MGE remains constrained by several practical limitations including prolonged incubation times (2–3 days), cytotoxicity at high precursor concentrations, and potential cellular stress due to altered glycosylation patterns.[12] These drawbacks limit its suitability for rapid, efficient, and broadly applicable membrane modifications.

Beyond genetic engineering and MGE, traditional bioconjugation strategies offer alternative routes for cell surface modification by chemically targeting lysine and cysteine residues.[13,14] However, these methods present their own challenges. Lysine residues, abundant on cell surfaces, carry a positive charge at physiological pH, and their modification can alter membrane charge and disrupt electrostatic interactions. Cysteine residues, though less abundant, are critical for maintaining protein structure through disulfide bonds; their modification risks destabilizing protein conformation, particularly in the oxidizing extracellular environment. For these reasons, we shifted our focus to the chemical modification of tyrosine residues. Tyrosines are neutral, and their modification does not disrupt the membrane’s overall charge, preserving the cell’s biochemical integrity. In this context, we previously developed click-electrochemistry (eY-click), a novel strategy for bioconjugating tyrosine residues on cell surfaces. This method offers rapid reaction kinetics and precise, time-dependent functionalization, enabling controlled labeling of tyrosine while minimizing nonspecific binding and structural disruption.[15]

Despite its many advantages, eY click requires electrochemistry equipment (electrodes, potentiostat). To explore a complementary approach, we developed a purely chemical alternative using aryl diazonium salt derivatives to modify tyrosine residues on cell surfaces (**Figure 1**).

Although diazonium-based tyrosine bioconjugation has been demonstrated in specific biological contexts, such as the modification of bacteriophage MS2 capsids [16,17] and our own studies on adeno-associated viruses,[18] its application for broader cell surface modifications remains underexplored. Ancient studies used diazonium salts for surface modification but did not benefit from advancements in molecular targeting and biomedical applications.[19–21] This highlights a gap in the field and the need for updated methods that leverage diazonium salts for broader, application-driven cell membrane modifications.

Here, we demonstrate that a diazonium salt-based strategy enables the rapid and efficient functionalization of diverse cell types, including immortalized cell lines and primary immune cells. This approach provides controlled, tyrosine-selective modification while circumventing the limitations commonly associated with metabolic glycoengineering (MGE) or lysine- and cysteine-targeted chemistries. Notably, we successfully modify both antigen-presenting and effector immune cells, thereby enabling interventions at multiple levels of the immune response. This strategy preserves cellular integrity across all tested cell types and offers significant potential for applications that demand precise and efficient cell surface engineering, such as targeted drug delivery, advanced diagnostics, and the development of therapeutic cell-based products. Furthermore, our versatile strategy enables ultrafast chemical modification, in one or two steps, of adherent, suspension, or primary cells for the chemical coupling of diverse molecules including fluorophores, carbohydrates, and biotin, as well as biomolecules like the Nanofitin scaffold.[22,23]

In summary, our strategy represents a significant advancement in cell surface engineering, paving the way for targeted biomedical applications, particularly in cancer therapy and vaccine development.

## 2. Results and discussion

The development of a universal methodology for chemically modifying cell membranes represents a potentially transformative advancement. Such an approach holds great promise as a versatile tool for researchers in both academic and private sectors, providing a robust and adaptable framework for mechanistic and therapeutic investigations across diverse cellular contexts. To this end, we synthesized the aryl diazonium salt **1** (**Figure 2-A**), as previously reported by our group. [24] This compound reacts *via* electrophilic aromatic substitution with the phenol side chain of tyrosine, forming a stable diazo bond. It was evaluated for its efficacy in bioconjugating tyrosine residues on membrane proteins in both adherent HeLa and suspension EXPI-f cells. These cell lines play a pivotal role in biomedical research, serving as essential models for exploring various aspects of cell biology, disease mechanisms, and innovative therapeutic strategies.

**Figure 2:**
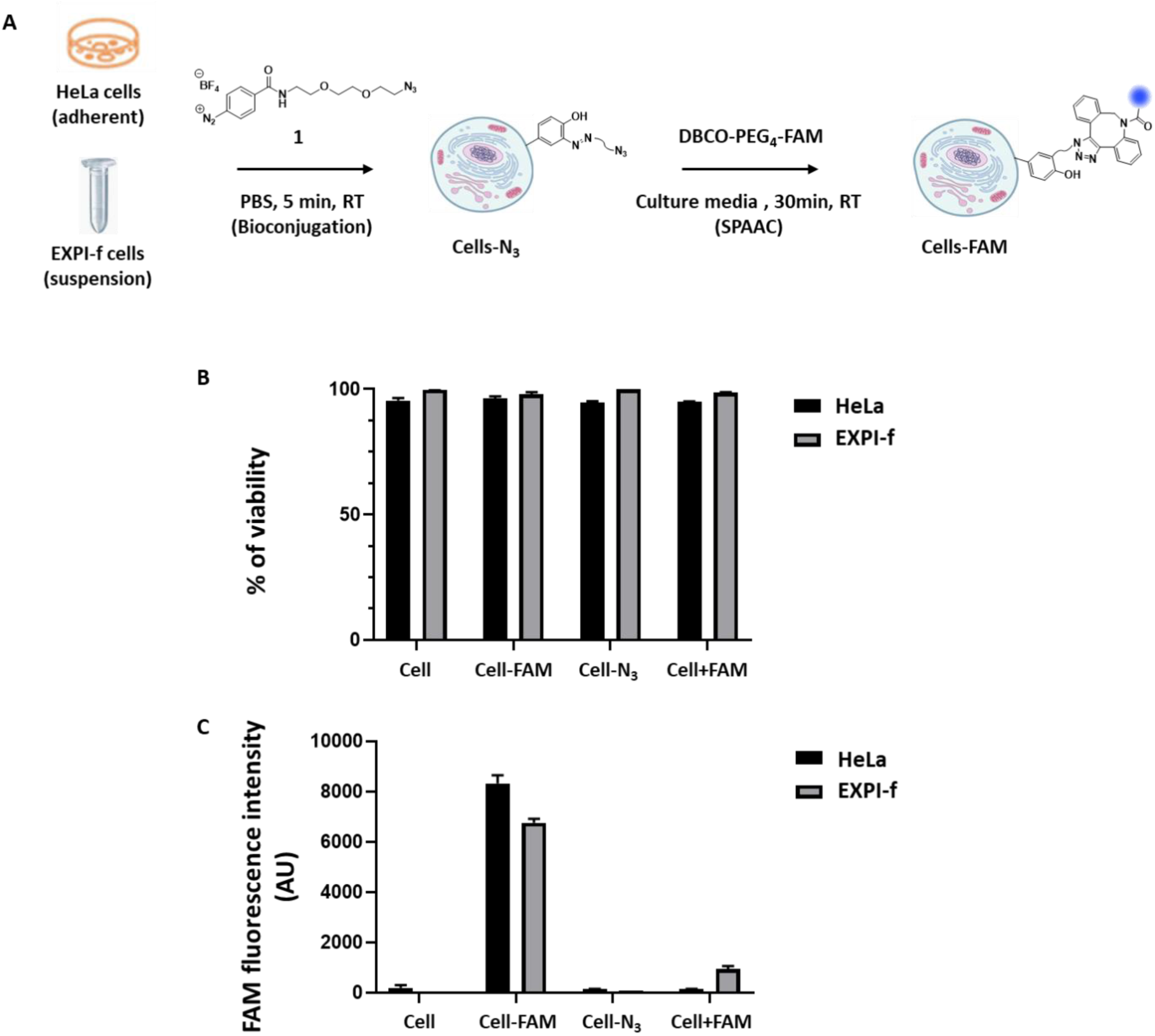
HeLa and EXPI-f cell membrane modification. **A)** HeLa cells and EXPI-f were conjugated 5 min with **1**, followed by 30 min with DBCO-PEG_4_-FAM (FAM). **B)** Viability % of unmodified HeLa and EXPI-f cells, HeLa-FAM and EXPI-f-FAM cells (modified with **1** (100 µM) and incubated with DBCO-PEG_4_-FAM (100 µM)), HeLa-N_3_ and EXPI-f-N_3_ cells (modified with **1** (100 µM)), and HeLa + FAM and EXPI-f + FAM cells (incubated only with DBCO-PEG_4_-FAM (100 µM)), as evaluated by flow cytometry after incubation with LIVE/DEAD® assay kit (*n* = 3). **C)** FAM fluorescence intensity in arbitrary unit (AU) measured by flow cytometry for the same samples (*n* = 3).

After solubilizing compound **1** in PBS buffer at a concentration of 100 µM, the solution was directly added to HeLa cells in the wells of the culture plate or to EXPI-f cells in a low-binding vial. Following a 5-minute incubation with gentle stirring, the PBS solution was carefully removed through either aspiration or centrifugation, repeated three times to eliminate excess reagent (**Figure 2-A**).

This first step enables the covalent attachment of the azide compound **1** to tyrosine residues on membrane proteins via electrophilic aromatic substitution, providing a bioorthogonal chemical handle for subsequent conjugation. In the second step, cells–N₃ were incubated for 30 minutes in the appropriate culture medium (DMEM for HeLa and BalanCD for EXPI-f cells) with 100 µM of the fluorescent reporter DBCO-PEG_4_-FAM (FAM), enabling strain-promoted azide-alkyne cycloaddition (SPAAC). This copper-free click reaction proceeds rapidly and selectively under physiological conditions, forming a stable triazole linkage without requiring metal catalysis, and is particularly well suited for live-cell applications.

To assess potential toxicity associated with the bioconjugation process, the cells viability was evaluated two hours post-grafting using the LIVE/DEAD® Viability/Cytotoxicity assay kit. As shown in **Figure 2-B**, the bioconjugation process did not induce any detectable toxicity in either HeLa or EXPI-f cells. Across all evaluated conditions, cells treated sequentially with compound **1** and DBCO-PEG_4_-FAM (Cell-FAM), cells treated with compound **1** alone (Cell-N_3_), and cells treated with DBCO-PEG_4_-FAM alone (Cell+FAM), more than 95% of the cells remained viable. All cells exposed to DBCO-PEG_4_-FAM exhibited fluorescence, regardless of prior bioconjugation, likely reflecting nonspecific adsorption or probe internalization. However, the relative fluorescence intensity of Cell-FAM (in arbitrary units) reached approximately 8000 for HeLa cells and 6000 for EXPI-f cells, whereas it remained around 100 and below 1000, respectively, in cells incubated with DBCO-PEG_4_-FAM alone (**Figure 2-C**, (gating strategy **Figure S1**). This pronounced contrast validates the efficiency of both the initial bioconjugation and subsequent SPAAC reaction, underscoring the robustness and specificity of our approach across adherent and suspension cell types. Taken together, these findings establish a powerful and non-toxic platform for cell membrane engineering applicable to a broad range of experimental settings.

Concurrent with the validation of our approach on conventional cell lines, we explored its feasibility on key immune cells, including antigen-presenting cells such as dendritic cells (DCs) and B cells, as well as effector cells such as T, B and NK cells. These immune cells represent a more clinically relevant and sophisticated model for testing bioconjugation technologies, paving the way for their potential application in advanced therapeutic contexts.[25] Dendritic cells, as the primary initiators of immune responses, play a pivotal role in antigen presentation and the activation of B and T lymphocytes.[26]. In addition to DCs, B cells can also act as antigen-presenting cells by displaying antigens on their MHC class II molecules to CD4⁺ T cells. This interaction induces B-cell activation and subsequent differentiation into plasma cells, leading to the secretion of antigen-specific antibodies.[27] T lymphocytes are key components of the adaptive immune cells, recognizing antigens with high specificity through their TCRs to eliminate infected or malignant cells and orchestrate immune responses.[28] NK cells, in contrast, are innate immune effectors that target virus-infected and tumor cells by sensing altered or missing MHC class I molecules, playing a pivotal role in tumor immunosurveillance.[29]

First, we compared the performance of the MGE strategy with our bioconjugation approach using DC2.4 cell line, which are adherent, thereby simplifying handling and analysis. Given that MGE has been reported to induce toxicity in A549 cells at concentrations exceeding 100 µM, [12] we first evaluated this parameter in DC2.4 cells using the LIVE/DEAD® Viability/Cytotoxicity assay kit prior to comparing the two methods (**Figure S2-A**). Consistent with the findings of Tomas *et al.* in A549 cells,[12] our results demonstrated that Ac_4_ManNAz concentrations exceeding 100 µM (^MGE^DC2.4-FAM) induced significant toxicity in DC2.4 cells, reducing cell viability to 30% at 250 µM (**Figure S2-B**). Furthermore, we confirmed the efficacy of the MGE approach for chemically modifying the DC2.4 cell membrane with an increase in fluorescence intensity observed following incubation with DBCO-PEG_4_-FAM. (**Figure S2-C**). Next, we performed the same experiment with DC2.4 cells, this time maintained in suspension (*i.e*. after trypsinization and before replating onto adherent surface), and similarly observed no toxicity. The fluorescence intensity resulting from the SPAAC reaction after incubation with Ac_4_ManNAz at 100 µM was similar to that obtained with adherent DC2.4 cells (**Figure S3**). Thus, for both suspension and adherent cells, 100 µM of Ac_4_ManNAz represents the maximum concentration that preserves cell viability.

We next compared the performance of MGE technology with our tyrosine bioconjugation strategy for membrane proteins in DC2.4 cells, under adherent and suspension culture conditions. For MGE, adherent DC2.4 cells were incubated in 24-well plates for 72 hours with Ac_4_ManNAz at 100 µM, while for the bioconjugation step, DC2.4 cells were exposed to compound **1** at concentrations of 100 µM and 250 µM for 5 minutes. Following the SPAAC reaction with the DBCO-PEG_4_-FAM (^MGE^DC2.4-FAM and DC2.4-FAM) probe at 100µM, no toxicity was observed under any conditions (**Figure 3-A**). For adherent cells, at 100 µM of compound **1**, the fluorescence intensity was comparable between the MGE and bioconjugation strategies, whereas at 250 µM, a slight increase was observed. This effect was amplified in suspension cells, where fluorescence intensity increased threefold at 250 µM of compound **1** (**Figure 3-B**), likely due to enhanced accessibility and a higher number of available tyrosine residues on membrane proteins.

**Figure 3:**
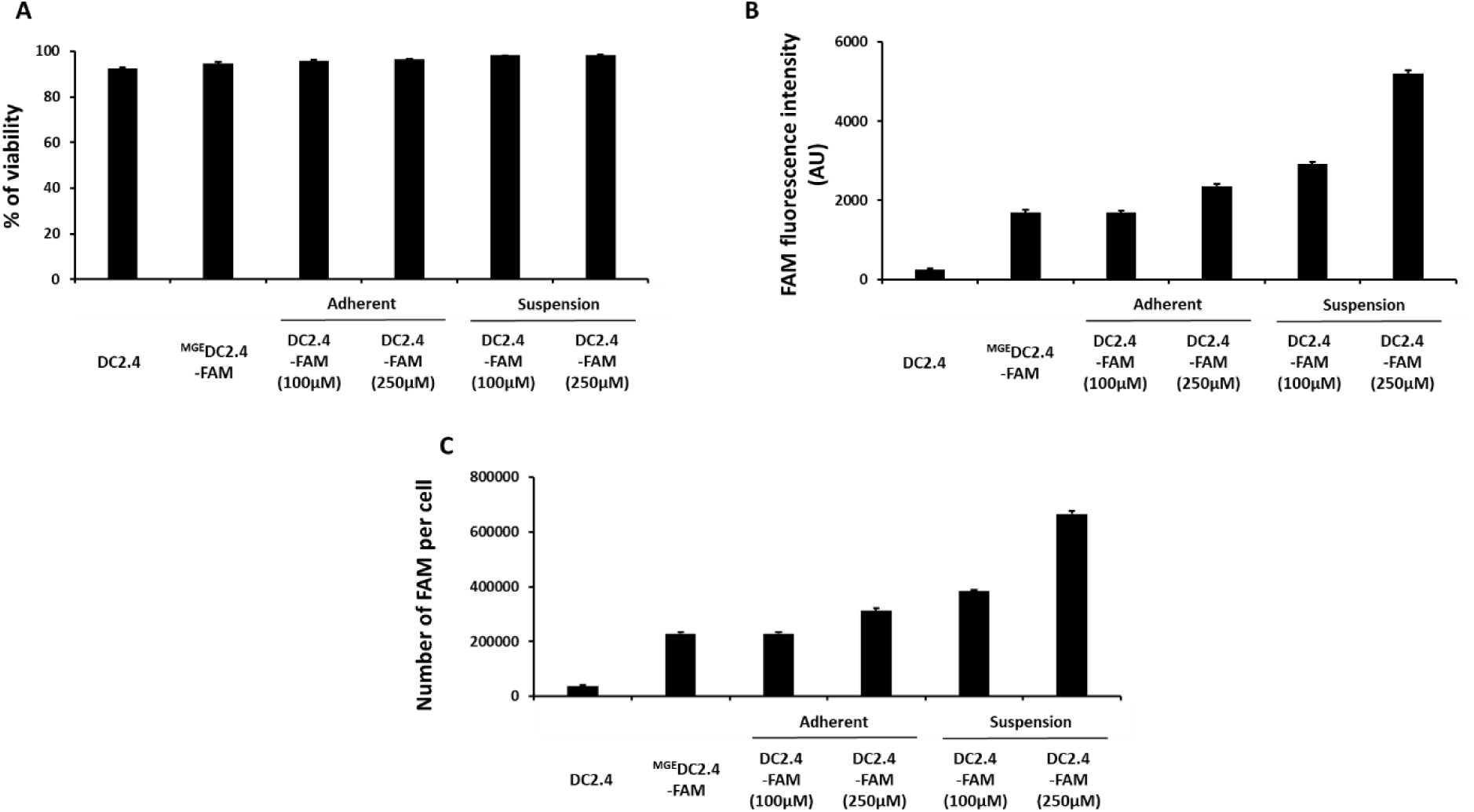
Comparison of MGE *versus* diazonium salt strategies. **A)** Viability % of unmodified DC2.4 cells, DC2.4 cells incubated with Ac_4_ManNaz at 100 µM for 72h, followed by 30 min SPAAC with DBCO-PEG_4_-FAM (100 µM, ^MGE^DC2.4-FAM), DC2.4-FAM adherent cells (modified with **1** (100 and 250µM)) and incubated with DBCO-PEG_4_-FAM (100 µM), and DC2.4-FAM suspension cells (modified with **1** (100 and 250µM) and incubated with DBCO-PEG_4_-FAM (100 µM) as evaluated by flow cytometry after incubation with LIVE/DEAD® assay kit (*n* = 3). **B**) FAM fluorescence intensity in arbitrary unit (AU) measured by flow cytometry for the six same samples (*n* = 3). **C)** Number of FAM per cell as evaluated by flow cytometry with the commercial Quantum^TM^ FITC kit for the six same samples (*n* = 3).

To quantify the number of FAM probes conjugated to the cell surface, we used the Quantum™ FITC kit, which enables precise calibration through five populations of FITC-labeled microspheres. Thus, with MGE, approximately 227,000 FAM were conjugated, whereas with bioconjugation, the highest concentration results in the grafting of 666,000 molecules per cell (**Figure 3-C**). Our approach allows for the modulation of ligand density on the surface of membrane proteins to a level that, to our knowledge, has not been previously reported. This tunability, unmatched by MGE technology, could be particularly valuable in vaccination models, immunotherapy, and other cell-based therapies, enabling the generation of chemically tailored cells for specific targets.

We next investigated the potential impact of bioconjugation on cell proliferation using the CellTrace™ Far Red kit, a fluorescent dye commonly used in flow cytometry to monitor cell division. Following the coupling of compound **1**, DC2.4-N_3_ cells were incubated with CellTrace™ Far Red and cultured under standard conditions. The SPAAC reaction with DBCO-PEG_4_-FAM was performed on day 0 to establish baseline fluorescence intensity prior to cell division, and was repeated after 24 and 48 hours of culture.

In unmodified DC2.4 cells, CellTrace™ fluorescence intensity decreased from approximately 60,000 at baseline to 7,000 after 24 hours and further to 2,000 at 48 hours (**Figure S4-A**). A comparable decay pattern was observed in DC2.4-N_3_-modified cells, with similar intensity values (**Figure S4-B**), indicating that the bioconjugation process does not interfere with cell proliferation. Analysis of fluorescence resulting from the SPAAC reaction showed a decrease in intensity following a similar trend (**Figure 4**). Quantification of bioconjugated FAM molecules using the Quantum™ FITC kit revealed a reduction of approximately 75% after 24 hours, followed by an additional 50% decrease after 48 hours. This reduction correlates with the CellTrace™ data, where the fluorescence reduction is more pronounced between 0 to 24 hours (**Figure 4**). This correlation demonstrates that bioconjugation preserves the cells’ physiological proliferative capacity. Notably, over 100,000 N_3_ molecules remained on the cell surface after 48 hours, underscoring both the stability of the modification and its potential for downstream applications.

**Figure 4:**
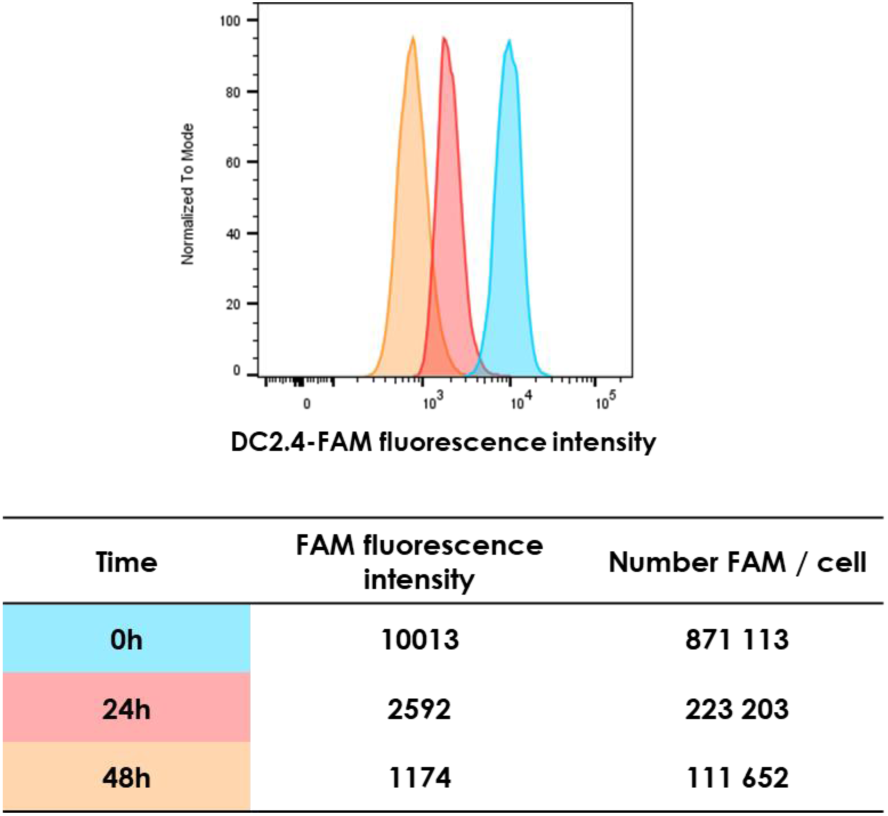
DC 2.4 cells proliferation. DC2.4-N_3_ cells (modified with **1** (100 µM)) were incubated with DBCO-PEG_4_-FAM (100 µM) (DC2.4-FAM) for the SPAAC reaction after 0h, 24h and 48h of cell culture and FAM fluorescence intensity in arbitrary unit (AU) measured by flow cytometry and number of FAM per cell as evaluated by flow cytometry with the commercial Quantum^TM^ FITC kit fluorescence (*n* = 3).

Importantly, we show that the bioconjugated signal diminishes over time due to cell division, offering a self-limiting alternative to permanent genetic modifications, such as CAR-T cells, thereby potentially mitigating risks associated with prolonged immune activation. Unlike CAR-T therapies whose long-term persistence can lead to chronic immune activation, and in rare cases, secondary malignancies due to insertional mutagenesis,[30] our approach inherently limits the duration of cellular functionalization through natural dilution, enhancing safety while preserving efficacy.

Given the importance of preserving and transporting cells while retaining the ability to perform SPAAC reactions after storage, we generated DC2.4-N_3_ cells using compound **1** at 100 µM, as previously described, and stored them at -80°C for 48 hours. Upon thawing, click-chemistry with DBCO-PEG_4_-FAM was performed for 30 minutes at 100 µM in culture medium, resulting in clear fluorescent labeling without any detectable cell mortality, confirming that the stored azide-functionalized cells retained their reactivity (**Figure S5**). These findings confirmed that DC2.4-N_3_ cells can be successfully frozen without any detrimental effects on the chemical modification process.

After validating the modification of HeLa, EXPI-f, and DC2.4 cell lines, we extended this strategy to B lymphocytes (B-EBV), T lymphocytes (Jurkat), and NK92 cell lines. As previously observed, no toxicity was detected, and fluorescence intensity was only observed when both components of the click chemistry reaction were present (**Figure S6**). These results underscore the broad applicability of our approach, enabling the targeted modification of tyrosine residues on cell membrane proteins and demonstrating its effectiveness across various cell types.

To further expand the scope of our work, we investigated the potential of our technology to bioconjugate larger biomolecules, such as proteins, that could enhance or modulate cellular functions. We generated a DBCO-conjugated Nanofitin and evaluated its ability to be grafted onto DC2.4 and NK92 cell lines. The Nanofitin used in this study carried a histidine-tag at its *N*-terminal end to facilitate detection, and was designed to target the epidermal growth factor receptor EGFR,[22,23] a receptor commonly overexpressed on cancer cell membranes.[31] Nanofitins present the advantage of having their *N*- and *C*-terminal ends oriented opposite to the binding site, allowing chemical modification at these termini without interfering with their molecular recognition properties. For anti-EGFR Nanofitin derivatization with DBCO, a cysteine residue was introduced at the *C*-terminal end, followed by thiol-based Michael addition with a maleimide-PEG_4_-DBCO derivative.

Following bioconjugation of DC2.4 and NK92 cells with compound **1** at 100 µM, the SPAAC reaction was performed at 37°C with 50 µM of DBCO-Nanofitin. After washing, an Alexa-488 anti-histidine tag antibody was used to detect Nanofitin on the cell surface via flow cytometry (DC2.4-NanoF or NK92-NanoF). To validate bioconjugation, we included the control DC2.4 or NK92 cells + DBCO-Nanofitin + Alexa-488 anti-histidine tag antibody (Ctrl).

Flow cytometry analysis revealed that 98% of DC2.4-NanoF cells and 80% of NK92-NanoF cells exhibited fluorescence (**Figure 5-A,B,C,D**), compared to a maximum of 5% under control conditions. Furthermore, fluorescence intensity reached approximately 500, higher than the baseline signal, confirming the efficient conjugation of biomolecules onto the cell membrane (**Figure 5-E,F**). The SPAAC click chemistry enables efficient bioconjugation of larger molecules such as proteins and, as in previous experiments, no toxicity was observed under these conditions (**Figure S7**).

**Figure 5:**
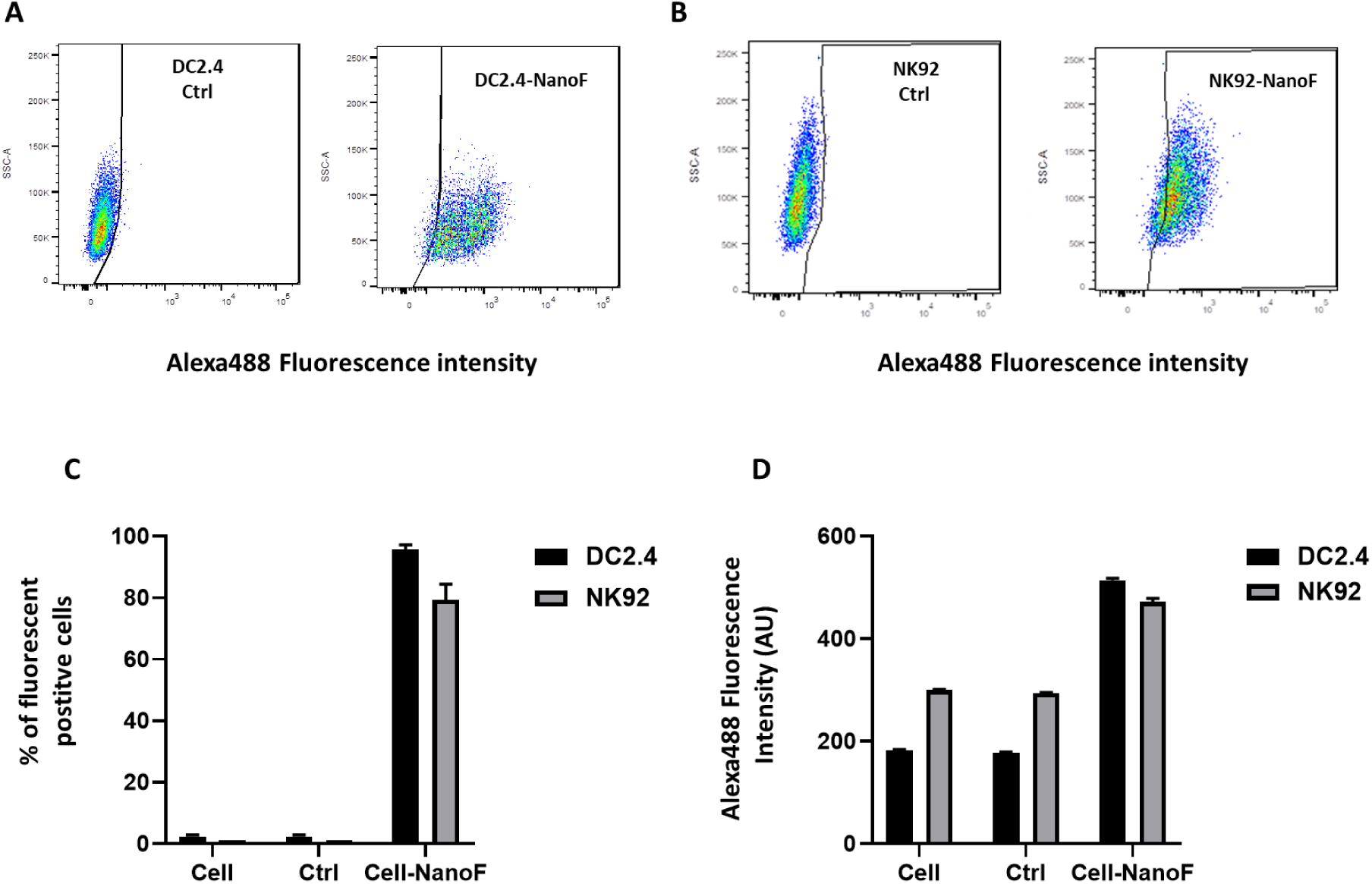
Chemical coupling of Nanofitin on Y of DC2.4 and NK92 protein cell membrane. **A,B**) Flow cytometry plots of Alexa488 fluorescence intensity of DC2.4 and NK92 cells incubated with DBCO-Nanofitin and Alexa-488 anti-histidine tag antibody (Ctrl) and DC2.4-NanoF and NK92-NanoF cells (modified with **1** (100 µM), incubated with DBCO-Nanofitin (50 µM)) and incubated with Alexa-488 anti-histidine tag antibody. **C**) % of fluorescent positive cell of non-modified DC2.4 and NK92 cells (Cell), DC2.4 and NK92 cells incubated with DBCO-Nanofitin and Alexa-488 anti-histidine tag antibody (Ctrl), DC2.4-NanoF and NK92-NanoF cells (modified with **1** (100 µM), incubated with DBCO-Nanofitin (50 µM)) (Cell-NanoF) and incubated with Alexa-488 anti-histidine tag antibody, as evaluated by flow cytometry (n=3). **D**) Fluorescence intensity of all the same three samples as evaluated by flow cytometry (n=3).

We demonstrated that the two-step approach, bioconjugation followed by the SPAAC reaction, enables the grafting of a wide range of structures, from fluorophores to proteins, underscoring its broad applicability. In addition, for non-peptidic or non-protein ligands, we evaluated the feasibility of a simplified one-step bioconjugation strategy. This approach bypasses the click-chemistry step and enables direct membrane functionalization within only 5 minutes, thereby reducing the overall process time from 1 hour to 5 minutes. To this end, we synthesized and characterized the biotin-N_2_^+^ derivative **2** through three steps, starting from the biotin amine (**Figure S8-13**), leveraging the strong biotin-streptavidin interaction, which is widely used in biological applications (**Figure 6-A**).[32] We also employed a GalNAc-N_2_^+^ derivative **3**, previously reported by our group (**Figure 6-A**),[18] selected for its potential to enhance cell recognition and anticancer properties, particularly against hepatocellular carcinoma, which overexpresses the asialoglycoprotein receptor (ASGPr).[33]

**Figure 6:**
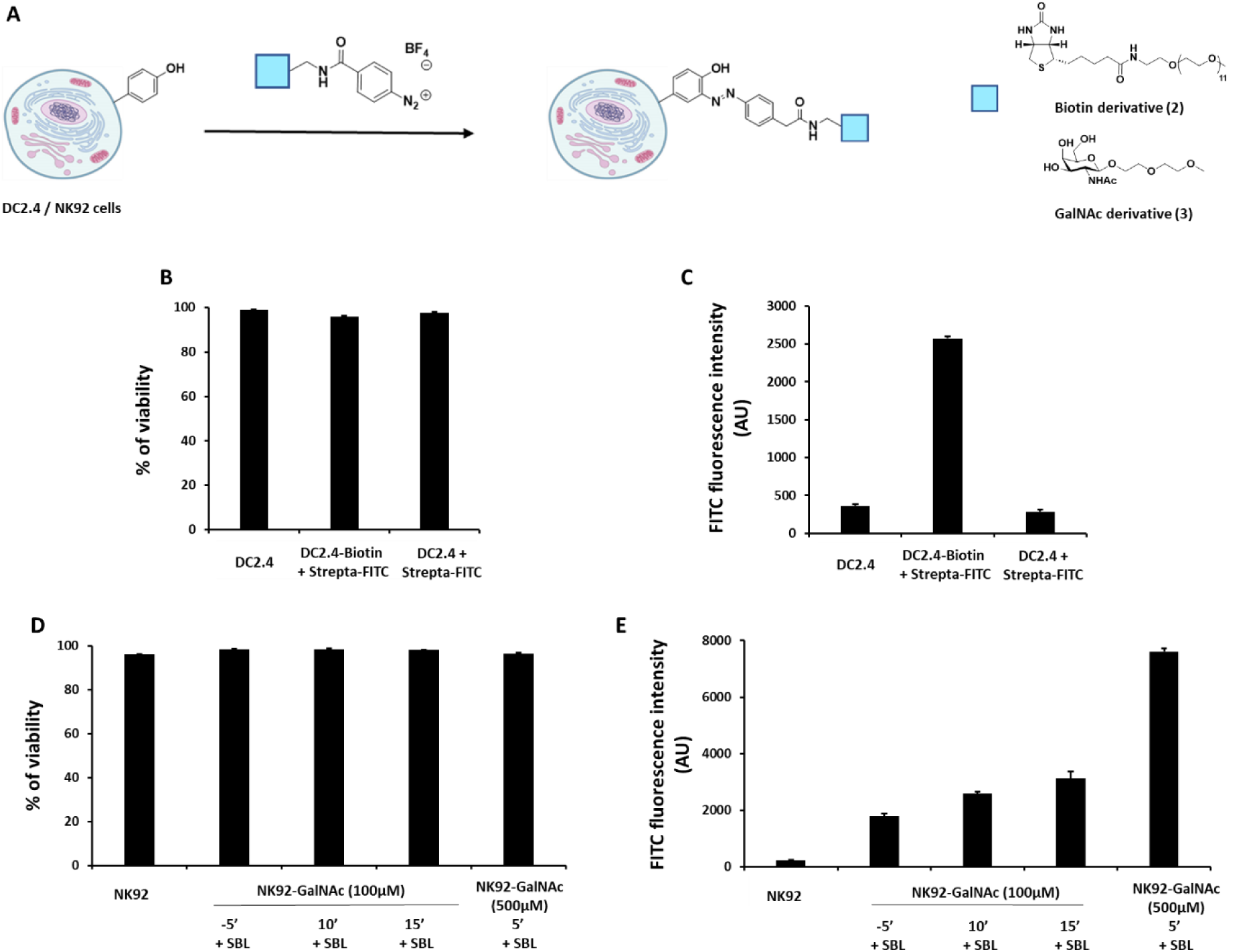
One-step bioconjugation of Y on protein cell membrane using a biotin and a GalNac diazonium salt derivative. **A)** DC2.4 and NK92 cells were conjugated 5 min with **2** or **3**. **B**) Viability % of unmodified DC2.4 cells, DC2.4 cells bioconjugated with **2** and incubated with streptavidin-FITC, DC2.4 cells incubated with streptavidin-FITC as evaluated by flow cytometry after incubation with LIVE/DEAD® assay kit (*n* = 3). **C**) FITC fluorescence intensity in arbitrary unit (AU) measured by flow cytometry (*n* = 3). **D**) Viability % of unmodified NK92 cells, NK92 cells bioconjugated with **3** at 100µM during 5’, 10’ and 15’, and at 500µM during 5’ and incubated with Soybean-FITC lectin (SBL) as evaluated by flow cytometry after incubation with LIVE/DEAD® assay kit (*n* = 3). **E**) FITC fluorescence intensity in arbitrary unit (AU) of the six same samples measured by flow cytometry (*n* = 3).

In this one-step approach, the biotin derivative **2** was covalently grafted onto tyrosine residues of cell surface proteins at a concentration of 100 µM on DC2.4 cells in suspension, yielding DC2.4-Biotin. These cells could serve as a basis for vaccination strategies involving engineered streptavidin-based antigen presentation. Following a 5-minute reaction in PBS buffer at room temperature and subsequent washing steps, no alteration in cell viability was observed. Flow cytometry analyses revealed a clear fluorescence signal upon staining with FITC-labeled streptavidin, confirming successful conjugation on the cell surface (**Figure 6-B, C**).

Similarly, the GalNAc derivative **3** was covalently attached to tyrosine residues on the surface of NK92 cells under different conditions: 100 µM for 5, 10, or 15 minutes, or 500 µM for 5 minutes in PBS buffer. GalNAc-modified NK92 cells may facilitate selective targeting of hepatocytes or hepatocellular carcinoma, given the high expression of ASGPr on liver cells as already described.[34] After washing, NK92-GalNAc cells were incubated with FITC-labeled soybean lectin to detect GalNAc residues on the cell membrane. No toxicity was observed under any of the tested conditions. Although a baseline fluorescence signal is expected due to the natural presence of GalNAc on the cell membrane, fluorescence intensity increased fourfold when 500 µM of compound **3** was used for 5 minutes (**Figure 6-D,E**). These results confirm both the successful grafting of exogenous GalNAc and the tunability of the system by modulating coupling conditions.

While the two-step strategy remains essential for the conjugation of complex biomolecules, such as proteins, the one-step method enables rapid and efficient functionalization of small molecules in just 5 minutes.

Together, these complementary approaches provide a versatile platform for tailoring cell surface properties through the grafting of a wide range of molecules according to structural complexity and intended application.

While the efficiency and non-toxicity of the bioconjugation process have been validated, it remains essential to ensure that chemically modified cells do not undergo activation. Cellular activation could trigger signaling pathways that alter their natural behavior, potentially compromising biological integrity and leading to biased or unreliable results. Moreover, activated cells may release cytokines or other mediators, potentially inducing inflammation or interfering with surrounding cellular processes.[35,36] Therefore, confirming that bioconjugation does not inadvertently activate cells is critical for maintaining both physiological relevance and safety in *in vitro* and *in vivo* applications.

To assess potential cellular activation following bioconjugation, we analyzed Jurkat and DC2.4 cells bioconjugated with compound **1** (Jurkat-N_3_ and DC2.4-N_3_). These cells were either left unstimulated or activated using PMA/ionomycin (P/I) for Jurkat cells, or lipopolysaccharides (LPS) for DC2.4 cells both for 24 hours. Activation markers were assessed by incubating Jurkat cells with anti-CD25 and anti-CD69 antibodies, and DC2.4 cells with anti-CD40, anti-CD86, and anti-MHC II antibodies.

In Jurkat cells, P/I stimulation led to a substantial increase in CD25 and CD69 expression, from approximately 1% to 90%, regardless of whether the cells had undergone bioconjugation, as determined by flow cytometry. CD25 (the IL-2 receptor α chain) and CD69 are early markers of T cell activation, indicative of functional immune responsiveness. In the absence of stimulation, no significant changes in activation marker expression were observed immediately after bioconjugation or following 24 hours of culture (**Figure 7-A**).

**Figure 7:**
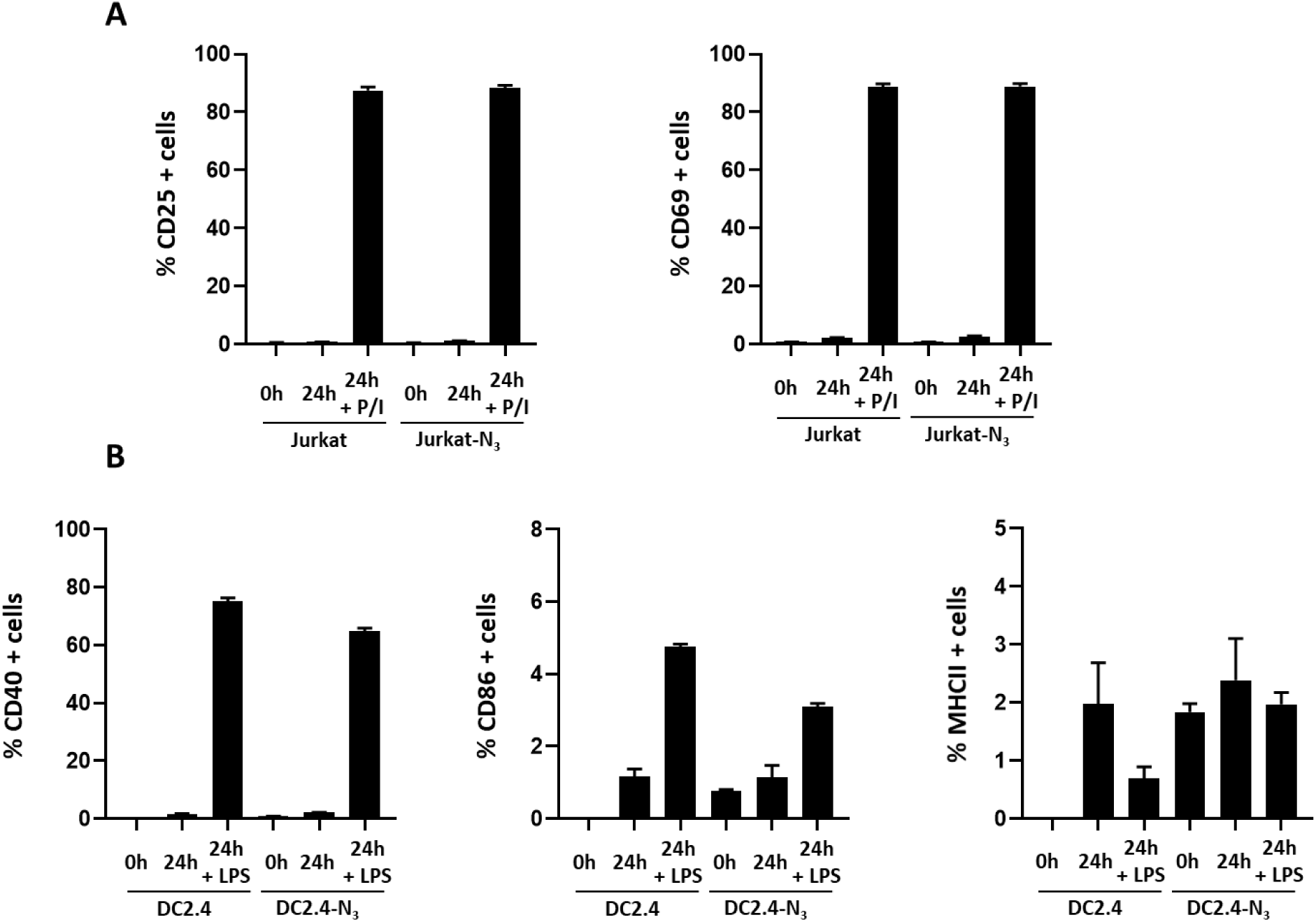
Activation profile of chemically modified Jurkat and DC2.4 cells. **A)** Percentage of CD25 and CD69 positive Jurkat and Jurkat-N_3_ cells (modified with **1** (100 µM)), without or with PMA-Ionomycin (P/I) activation, were incubated with anti-CD25 and anti-CD69 fluorescent antibody, as evaluated by flow cytometry (n=3). B) Percentage of CD40, CD86 and MHCII positive DC2.4 and DC2.4-N_3_ cells (modified with **1** (100 µM)), without or with LPS activation, were incubated with anti-CD40, anti-CD86 and anti-MHCII fluorescent antibody, as evaluated by flow cytometry (n=3).

Similarly, both modified and unmodified DC2.4 cells exhibited strong activation after 24 hours of LPS treatment, with CD40 expression increasing from ∼1% to 70%, and CD86 from ∼0.5% to 3–4%. CD40 and CD86 are co-stimulatory molecules typically upregulated on antigen-presenting cells upon immune activation. MHC class II expression, a marker of antigen presentation capacity, remained unchanged under all conditions. Again, no differences were observed between bioconjugated and control cells, either immediately after modification or after 24 hours in culture (**Figure 7-B**).

These results confirm that cell membrane modification via bioconjugation with compound **1** does not induce unintended cellular activation. Importantly, the modified cells retained their ability to be activated upon appropriate stimulation. This finding underscores the safety and reliability of the bioconjugation process, ensuring that modified cells remain physiologically functional and suitable for further biological applications.

We then extended our study to a clinically relevant context by applying the bioconjugation strategy to Peripheral Blood Mononuclear Cells (PBMCs) obtained from healthy donors. Unlike immortalized cell lines, these primary cells retain the heterogeneity and intrinsic characteristics of their tissue of origin, providing a model that more closely mimics physiological conditions. This is essential for evaluating both the efficacy and safety of our approach under biologically relevant settings. Moreover, the efficient modification of primary cells highlights the potential of this technology for personalized therapeutic applications, where bioconjugation strategies could be adapted to individual patient profiles.

PBMCs comprise a diverse population of immune cells, including B and T lymphocytes, NK cells, and dendritic cells. PBMCs isolated from three voluntary blood donors were subjected to the previously established protocol: an initial coupling step with compound **1** (100 µM), followed by a 30-minute SPAAC reaction with DBCO-PEG_4_-FAM (100 µM). As in previous experiments, cell viability remained high (∼90%), confirming the non-toxic nature of the process (**Figure S14-A**). The overall fluorescence intensity of chemically modified PBMCs reached approximately 8000, compared to ∼1000 in cells treated with DBCO-PEG_4_-FAM alone and ∼100 in non-modified control (**Figure S14-B**). These findings confirm that our tyrosine-based bioconjugation strategy is both efficient and broadly applicable across diverse primary immune cell types. Importantly, the ability to chemically modify blood donor–derived PBMCs while maintaining high viability represents a significant step toward potential clinical translation.

Next, we functionalized primary NK cells using the EGFR-targeting Nanofitin, which had previously been validated on the DC2.4 and NK92 cell lines as model systems. Two NK subpopulations were used: (i) resting NK cells within PBMCs and (ii) activated and expanded NK cells derived from PBMCs (**Figure S15**). Grafting an EGFR-specific Nanofitin onto the NK cell surface was designed to provide a ligand capable of promoting a more stable immunological synapse with EGFR-positive tumor cells, thereby conferring target specificity and enhancing NK cell cytotoxicity. Cells from five independent donors were used for each group. Briefly, cells were incubated for 5 minutes with compound **1** (100 µM), followed by a 30-minute SPAAC reaction with DBCO-Nanofitin (50 µM). After washing, surface display of the EGFR-specific Nanofitin was confirmed by flow cytometry using an Alexa-488 anti-histidine tag antibody (Cell-NanoF). For Labeling specificity, we included the control resting NK cells and activated NK cells + DBCO-Nanofitin + Alexa-488 anti-histidine tag antibody (Ctrl).

Flow cytometry analysis revealed efficient membrane conjugation, with approximately 95% of activated NK-NanoF cells and 45% of resting NK-NanoF cells exhibiting strong fluorescence, compared to less than 5% in control conditions (**Figure 9-A**). No cytotoxicity was detected under these conditions, consistent with prior experiments.

**Figure 9:**
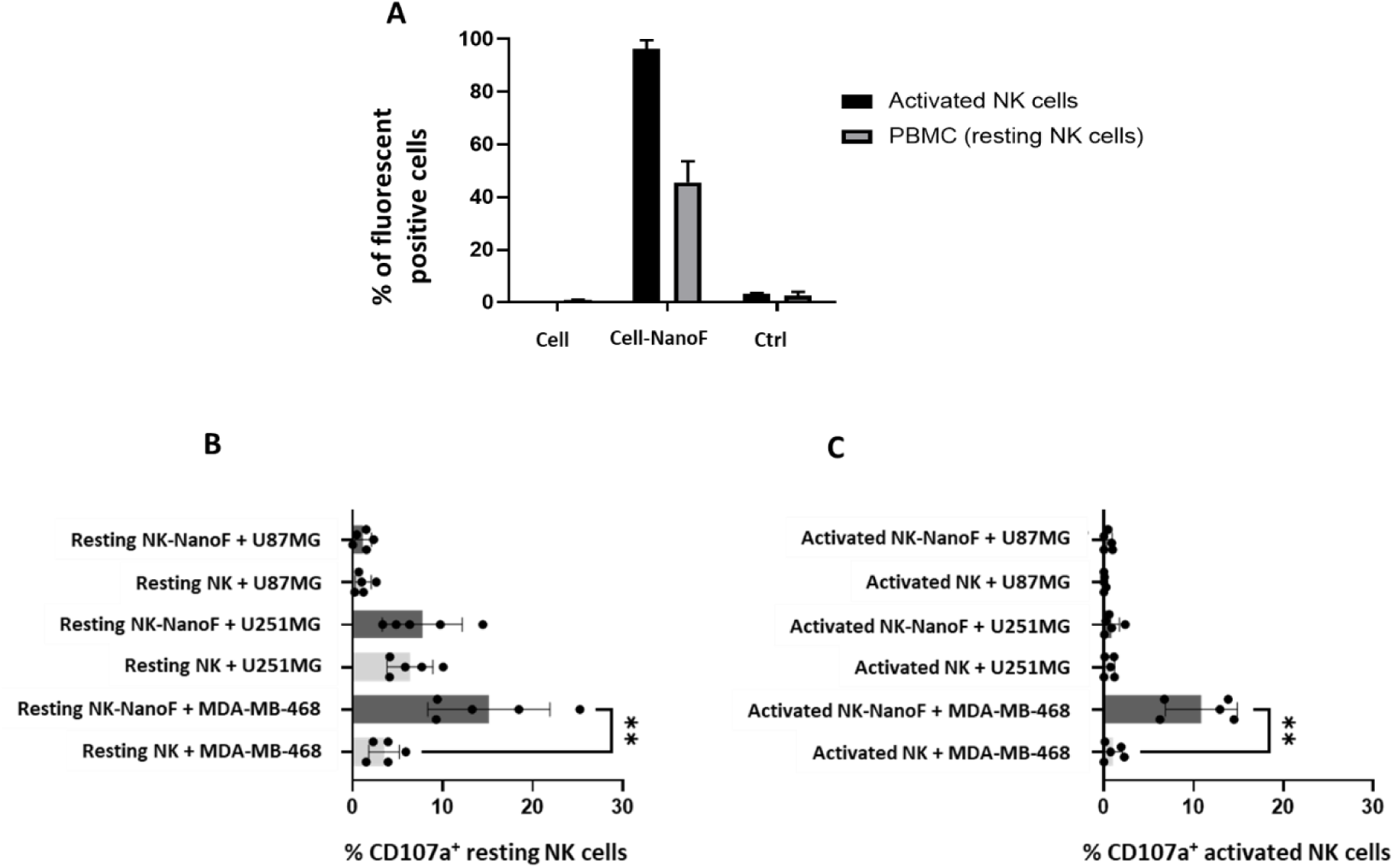
Chemical coupling of Nanofitin on Y of PBMC protein cell membrane. **A**) % of fluorescent positive cell of activated NK cells and resting NK cells, activated NK-NanoF and resting NK-NanoF (modified with **1** (100 µM), and incubated with DBCO-Nanofitin (50 µM)) and incubated with Alexa-488 anti-histidine tag antibody, activated NK and resting NK cells incubated with DBCO-Nanofitin (50 µM) and Alexa-488 anti-histidine tag antibody (Ctrl), as evaluated by flow cytometry (n=2). **B**) Bar graphs comparing degranulation of resting NK cells and resting NK-NanoF against U87MG, U251MG, and MDA-MB-468, after staining with anti-CD107a antibody, by flow cytometry (n=5). **C**) Bar graphs comparing degranulation of activated NK cells and activated NK-NanoF against U87MG, U251MG, and MDA-MB-468, after staining with anti-CD107a antibody, by flow cytometry (n=5). Statistical significance is indicated as follows: **<0.01. Comparisons were made by Wilcoxon test.

Prior to functional assays, EGFR surface expression was assessed on three tumor cell lines: U87MG, U251MG, and MDA-MB-468. Flow cytometry following incubation with a fluorescently labeled anti-EGFR antibody revealed high EGFR expression on MDA-MB-468 cells, moderate expression on U251MG cells, and no detectable expression on U87MG cells (**Figure S16**), in agreement with previously reported data for these cell lines. [37]

To evaluate cytotoxic function, resting and activated NK-NanoF cells were co-cultured with these tumor cell lines at an effector-to-target ratio of 1:1. Degranulation activity was measured after five hours by detecting CD107a (LAMP-1), a transient surface marker of NK cell cytotoxicity (**Figure S17**).

Functionally, membrane display of the EGFR-specific Nanofitin enhanced immune cell cytotoxicity in a target-dependent manner. For resting NK cells, CD107a expression increased from 4% to 15% after incubation with MDA-MB-468 cells, with no significant change observed for U251MG (∼8%) or U87MG (∼2%) targets, consistent with their EGFR expression levels (**Figure 9-B**). For activated NK-NanoF cells, CD107a expression increased from 2% to 12% when co-cultured with MDA-MB-468 cells, with no effect observed for U251MG or U87MG (**Figure 9-C**).

Together, these results confirm that membrane-anchored Nanofitin can effectively redirect NK cell activity toward EGFR-positive tumor cells without requiring permanent genetic modification. This strategy therefore represents a flexible, non-genetic, and transient approach for arming immune cells with tumor-targeting capabilities, offering a promising alternative or complement to CAR-based therapies.

## 3. Conclusion

In this study, we developed a novel one- and two-step bioconjugation strategy that enables rapid and efficient chemical modification of tyrosine residues on the cell surface via diazonium salt chemistry. This ultra-fast, robust, and versatile method was validated across a broad range of cell types, including adherent, suspension, and primary human cells, using diverse ligands ranging from small molecules to proteins such as Nanofitin. Importantly, membrane functionalization was achieved without inducing cytotoxicity or undesired activation of dendritic cells (DC2.4) or T lymphocytes (Jurkat), highlighting the biocompatibility of the method.

We also demonstrated that the bioconjugated signal progressively decreases over time due to membrane turnover and cell division, offering a self-limiting and reversible alternative to permanent genetic modifications such as CAR-T engineering. This feature may help mitigate long-term side effects or uncontrolled immune activation, improving the safety profile of cell-based therapies.

The successful surface engineering of blood donor-derived PBMCs underscores the translational potential of this approach. In particular, we showed that grafting an EGFR-targeting Nanofitin onto immune cells (activated and resting NK) enhances their cytotoxic response against EGFR-positive tumor cells. These findings suggest that our platform could serve as a non-genetic, modular, and transient alternative to CAR-based therapies, offering greater control, lower manufacturing complexity, and enhanced safety.

Beyond its biological implications, the operational flexibility of the two-step strategy paves the way for practical applications, such as the creation of azide-functionalized cell banks (e.g., frozen at −80 °C) and ready-to-use reagent kits for customizable surface labeling.

Future work will aim to expand the range of graftable ligands, improve *in vivo* stability and targeting, and assess therapeutic efficacy in relevant animal models. By combining speed, adaptability, and safety, this cellular platform represents a powerful new tool for both fundamental research and next-generation biomedical applications.

## 4. Experimental methods

### Materials

Most of chemical reagents were purchased from Sigma Aldrich, Carbosynth, TCI Chemical and were used without further purification. Solvents such as DCM, Toluene, DIPEA, Pyridine, MeOH, MeCN or DMF were purchased anhydrous from Sigma Aldrich. DBCO-PEG_4_-FAM was purchased from Conju-Probe. Biotin amine derivative was purchased from BLDPharm. All chemically synthesized compounds were characterized by ^1^H (400.133 or 300.135 MHz), ^13^C (125.773 or 75.480 MHz) NMR spectroscopy (Bruker Avance 300 Ultra Shield or Bruker Avance III 400 spectrometer). Chemical shifts are reported in parts per million (ppm); coupling constants are reported in units of Hertz [Hz]. The following abbreviations were used: s = singlet, d = doublet, t = triplet, q = quartet, quin = quintet, br = broad singlet. When needed, ^13^C heteronuclear HMQC and HMBC were used to unambiguously establish structures. High resolution mass spectra (HR-MS) were recorded on a Waters Xevo G2-XS Qtof spectrometer coupled to an Acquity H-class LC apparatus. The ionization sources were performed with the available methods (ESI+, ESI-, ASAP+, ASAP-). A tolerance of 5 ppm was applied between calculated and experimental values. Column chromatography was conducted on silica gel Kieselgel SI60 (40−63 μm) from Merck, or on Silica cartridge and eluted via a puriFlash 430 with an UV and ELSD detection.

FITC-Soybean lectin was purchased from Vector Laboratories. FITC-Streptavidin was purchased from Thermofisher. Alexa488 anti HisTag antibody was purchased from RetD systems. FITC Soybean lectin was purchased from Vector Laboratories. MHC Class II (I-A/I-E) Monoclonal Antibody (M5/114.15.2), Brilliant Violet 421 and CD11b Monoclonal Antibody (M1/70), Super Bright 600 were purchased from Invitrogen. PE/Cyanine7 anti-mouse CD40 antibody was purchased from Biolegend.

NK cell degranulation was assessed using anti-CD107a-BV510 (H4A3, Sony). NK cells were phenotyped as CD3-CD56+ using anti-CD3-APC-Cy7 (SK7, Sony) mAb and anti-CD56-PE (HCD56, Sony).

U87MG, U251MG and MDA-MB-468 cell lines phenotyping was assessed using anti-EGFR-PE (528, Santa Cruz sc-120) mAb.

Live/Dead™ Fixable viability dye was purchased from Invitrogen (Catalog number L34957). Quantum™ FITC-5 MESF was purchased from Biorad (Catalog number FCSC555).

Flow cytometry was done on BD LSRFortessa™ X-20 or BD FACSCanto™ II (Becton Dickinson).

### Synthesis

The synthetic steps leading to compound **2**, along with the corresponding ¹H and ¹³C NMR spectra, are provided in the Supporting Information section (**Figure S8-S13**).

### Cell lines, primary cells and culture

All the cell lines HeLa, EXPI-f, DC2.4, NK92, Jurkat and BEB-V are not recorded in the misidentified cell lines list. They were cultured based on ATCC protocol. The HeLa, EXPI-f, DC2.4N, NK92, Jurkat and BEB-V cell lines were kindly provided by academic laboratories and are regularly tested for mycoplasma free and sterility. U87MG, U251MG and MDA-MB-468 cell lines were used to investigate NK cell degranulation. U87MG and U251MG were kindly provided by Dr Claire Pecqueur (UMR-S 1307, Nantes, France) and MDA-MB-468 was provided by Dr Philippe Juin (UMR-S 1307, Nantes, France). Mycoplasma tests performed by PCR were negative for all cell lines. PBMCs were isolated as previously described.[38] All blood donors (n=10) were recruited at the Etablissement Français du Sang (EFS, Nantes, France), and informed consent was given by all individuals. A declaration of preparation and conservation of these biocollections (AC-2021-4397) has been provided to French Research Minister and has received approval from the IRB (2015-DC-1).

Hela cells were cultured in DMEM supplemented with 10% FBS serum and 1% penicillin-streptomycin at 37°C under 5% CO_2_ atmosphere. EXPI-f cells were cultured in BalanCD medium (Catalog ID 91165 Fujifilm) supplemented with 4mM l-glutamine at 37°C under 5% CO_2_ atmosphere. DC2.4 cells were cultured in RPMI 1640 (sigma R8758) supplemented with 1% penicillin-streptomycin, 1% of l-Glutamine, 1% of HEPES buffer 1M, 0.1% of 2-betamecaptoethanol 50mM, 1% of MEM NEA 100X (Life Technology, 11140-035), 1% of sodium pyruvate 100mM (Sigma S8636) and 10% of SVF at 37°C under 5% CO_2_ atmosphere. NK92 cells were cultured in RPMI 1640 (sigma R8758) supplemented with 1% penicillin-streptomycin, 1% of l-Glutamine, 10% of SVF and 200 U.I of IL2 at 37°C under 5% CO_2_ atmosphere. BEB-V cells were cultured in RPMI 1640 (sigma R8758) supplemented with 1% penicillin-streptomycin, 1% of l-Glutamine, and 10% of SVF at 37°C under 5% CO_2_ atmosphere. Jurkat cells were cultured in cultured in RPMI 1640 (sigma R8758) supplemented with 1% penicillin-streptomycin, and 10% of SVF at 37°C under 5% CO_2_ atmosphere.

U87MG, U251MG and MDA-MB-468 cell lines were cultured in DMEM high glucose containing glutamine (Gibco) and penicillin-streptomycin (Gibco) supplemented with 10% of fetal bovine serum (Gibco) at 37°C in a humidified atmosphere with 5% CO_2_.

Resting NK cells in PBMCs were cultured in RPMI 1640 medium (Gibco, Paisley, Scotland, UK) containing glutamine (Gibco) and penicillin-streptomycin (Gibco) and supplemented with 10% fetal bovine serum (Gibco) at 37°C in a humidified atmosphere with 5% CO_2_. Activated and expanded NK cells derived from PBMCs were cultured in NK MACS medium (Miltenyi Biotec) supplemented with 10% HS, 1% GlutaMAX and 200U/mL IL-2 at 37°C in a humidified atmosphere with 5% CO_2._

### DBCO-Nanofitin production

E. coli DH5α clones expressing his-tag Nanofitin B10 with a C-terminal cysteine was cultivated in 2YT medium, in shake-flasks (37°C, 200 rpm). Nanofitin expression was induced with IPTG (1 mmol/L) for 4 hours. Cells were harvested by centrifugation (15 minutes, 4000 g, 4°C) using a Sorvall BIOS 16. Biomass was disrupted in SinapTec homogenizer, and cell debris removed by centrifugation (30 minutes, 12 000 g, 4°C). Supernatants were clarified by filtration through a 0.2 μm filter. Filtrate was treated by IMAC (HisTrap HP column) and then concentrated (2 kDa MWCO). After purification, Nanofitin was polished by size exclusion chromatography using a Superdex 75 column (Cytiva). Purified Nanofitin was formulated in Sodium phosphate 100mM, NaCl 200mM, EDTA 5mM, pH6.5, and filter with Mustang E 0.2µm (Cytiva) for endotoxin removal and sterilization. Protein purity was addressed using standard SDS-PAGE analysis and mass spectrometry. Endotoxin levels were assessed using the Endosafe-PTS LAL analysis (Charles River).

Before conjugation, Nanofitin was reduced with 10 equivalent molar of TCEP at 4°C -ON. For conjugation, 13.5 equivalent molar of Maleimide-PEG4-Dibenzocyclooctyne (Mal-PEG4-DBCO) (Fisher Scientific) were added for 30 minutes. The conjugation reaction was stopped by added 4 equivalent molar of L-cysteine (Merck) and Nanofitin was then formulated in PBS pH7.4 using HiTrap desalting column (Cytiva).

Nanofitin conjugated with Mal-PEG4-DBCO was quantified with BCA assay and purity was evaluated with SDS-PAGE and reverse-phase chromatography on ACQUITY UPLC Peptide CSH C18 Column, 130Å, 1,7μm, 2.1mm x 100mm (Waters).

### Bioconjugation, SPAAC reaction, MGE and characterizations

3.10^6^ cells were seeded in P24 well plates or suspended in a low binding vial in 500 µL of PBS. Compounds **1**, **2** and **3** were freshly prepared and added directly to the cells at 100µM for **1** and **2** and 100 and 500µM for **3**. After incubation with gentle stirring during 5 minutes for **1** and **2**, and 5, 10 and 15 minutes at 100 µM and 5 minutes at 500µM for **3**, the PBS solution was carefully removed through either aspiration (adherent cells) or centrifugation (suspension cells), repeated three times to eliminate excess reagent. The GalNAc- and biotin-cells were directly characterized. The N_3_-cells were then resuspended 500 µL of their specific culture medium and DBCO-PEG_4_-FAM was added at 100µM and DBCO-Nanofitin at 50 µM. The SPAAC reaction was done during 30 minutes at RT for DBCO-PEG_4_-FAM and at 37°C for DBCO-Nanofitin, with gentle stirring. The culture medium was carefully removed through either aspiration (adherent cells) or centrifugation (suspension cells), repeated three times to eliminate excess reagent.

MGE of DC2.4 cell surface glycans was performed using peracetylated N-azidoacetylmannosamine (Ac_4_ManNAz). Cells were seeded at a density of 2.10^5^ in P24 well plates or suspended in a low binding vial and cultured under standard conditions in complete medium supplemented at 50, 100 and 250 µM of Ac_4_ManNAz for 72 hours. The medium was then carefully removed through either aspiration (adherent cells) or centrifugation (suspension cells), repeated three times to eliminate excess reagent. Culture medium was added to the ^MGE^DC2.4 cells followed by addition of DBCO-PEG_4_-FAM at 100µM. The SPAAC reaction was done during 30 minutes with gentle stirring. The culture medium was then carefully removed through either (adherent cells) or centrifugation (suspension cells), repeated three times to eliminate excess reagent.

For FAM flow cytometry analyses, after centrifugation cells were resuspended in PBS 1X, SVF 2%, EDTA 2mM. For GalNAc detection, FITC Soybean lectin solution was added to the cells and incubated at RT during 30 minutes. Cells were then centrifugated and resuspended in specific medium before flow cytometry analyses. For biotin detection, FITC streptavidin was added to cells and incubated at RT during 30 minutes. Cells were then centrifugated and resuspended in specific medium before flow cytometry analyses. For Nanofitin detection, Alexa488 anti HisTag antibody was added to the cells and incubated at 37°C during 30 minutes. Cells were then centrifugated and resuspended in specific medium before flow cytometry analyses.

### Live /Dead™ assay, Cell Trace™ assay and FAM quantification

Aqua LIVE/DEAD™ Fixable viability dye (Invitrogen, Catalog number L34957) stock solution was prepared at by adding the appropriate volume of DMSO. Modified and non-modified cells were resuspended in 200 µL of PBS/ LIVE/DEAD™ (dilution 1/1000) and then incubated at 4°C in the dark during 20 minutes. Cells were centrifugated and resuspended in specific medium before flow cytometry analyses. LIVE/DEAD™ was excited with a 405 nm laser and detected with the 525/50 bandpass filter.

CellTrace™ Far Red (Invitrogen, Catalog number C34564) stock solution was prepared at 5mM (as manufacturer instruction) by adding the appropriate volume of DMSO. CellTrace stock solution was used at 5µM final working concentration. Modified and non-modified cells were incubated with the CellTrace™ solution for 20 minutes at 37°C in the dark. A five-times volume of warmed complete medium was added to cells for 5 minutes incubation. Cells were centrifugated and resuspended in specific medium before flow cytometry analyses or seeding for culture. CellTrace™ was excited with a 642 nm laser and detected with the 670/30 bandpass filter.

Quantum™ FITC-5 MESF (Biorad, Catalog number FCSC555) was used to quantitate the FITC fluorescence intensity in Molecules of Equivalent Soluble Fluorochrome units. FAM and Quantum™ FITC-5 beads were excited with a 488 nm laser and detected with the 530/30 bandpass filter.

Datas were analysed using software Flowjo.

### Activation of DC2.4 and Jurkat cells

Modified and non-modified DC2.4 cells were activated with 2 µL of lipopolysaccharide solution (500x) in a P6 plate with 1mL of culture medium, then incubated for 24 hours at 37°C in a humidified atmosphere with 5% CO_2_ and finally labelled with anti-MHC Class II monoclonal antibody, anti-CD11b Monoclonal Antibody, anti CD40 PE/Cyanine7 anti-mouse anti CD86 and anti B7-1/CD80 Antibody. Cells were then centrifugated and resuspended in specific buffer before flow cytometry analyses.

Modified and non-modified Jurkat cells were activated with 40µL of Phorbol 12-myristate 13-acétate “PMA” (P8139-1MG, Sigma) pre-diluted 1/1000, and 200µL of Calcium Ionophore A23187 hemicalcium salt “IONO” (C9275-1MG, Sigma) pre-diluted 1/100 in a final volume of 2 mL of culture medium, then incubated for 24 hours at 37°C in a humidified atmosphere with 5% CO_2_ and finally labelled with anti-CD3 BUV395 antibody, anti-CD4 APHC7 antibody, anti-CD69 BV421 antibody and anti-CD25 RB780 antibody. Cells were then centrifugated and resuspended in specific buffer before flow cytometry analyses.

### Degranulation of NK cells

Resting NK cells, resting NK-NanoF cells, activated NK cells or activated NK-NanoF cells were incubated with target cell lines in the presence of anti-CD107a-BV510 (H4A3, Sony) mAb, and NK-cell degranulation was assessed after incubation for 5h with different target cells (E:T ratio = 1:1) or medium (negative control) with brefeldin A (Sigma) at 10 µg/mL for the last 4 h. Then, NK cells were phenotyped as CD3^-^CD56^+^ by flow cytometry on a FACSCanto II cytometer. Systematically before acquisition, BD FACSDiva CS&T Research Beads (BD Biosciences) were used. The flow cytometry files were analysed using FlowjoTM 10.6 software (LLC, Ashland, OR).

## CRediT authorship contribution statement

**Mohammed Bouzelha** (Conceptualization; Methodology; Formal analysis), **Karine Pavageau** (Conceptualization; Methodology; Formal analysis), **Sarah Renault** (Conceptualization; Methodology; Formal analysis), **Enora Ferron** (Conceptualization; Validation; Formal analysis; Methodology; Writing – review & editing), **Nicolas Jaulin** (Conceptualization; Methodology; Formal analysis), **Maia Marchand** (Methodology; Formal analysis), **Dimitri Alvarez-Dorta** (Conceptualization; Methodology; Validation; Formal analysis), **Roxanne Peumery** (Methodology), **Mathieu Scalabrini** (Methodology, Writing – review & editing), **Héloïse Delépée** (Methodology), **Allwyn Pereira** (Methodology; Writing – review & editing), **Elise Enouf** (Conceptualization; Validation; Formal analysis; Methodology), **Mathieu Cinier** (Conceptualization; Validation; Formal analysis; Methodology; Writing – review & editing), Oumeya Adjali, **Sébastien G. Gouin** (Conceptualization; Methodology; Writing – review & editing), **Christelle Retière** (Conceptualization; Validation; Formal analysis; Methodology; Writing – review & editing), **Mickaël Guilbaud** (Conceptualization; Validation; Formal analysis; Methodology; Writing – review & editing), **David Deniaud** (Conceptualization; Funding acquisition; Supervision; Validation; Writing – original draft; Writing – review & editing), and **Mathieu Mével** (Conceptualization; Funding acquisition; Supervision; Validation; Writing – original draft; Writing – review & editing).

## Acknowledgements

This research was supported by the Fondation d’Entreprise Thérapie Génique en Pays de Loire, the Centre Hospitalier Universitaire (CHU) of Nantes, the Institut National de la Santé et de la Recherche Médicale (INSERM), the Centre National de la Recherche Scientifique (CNRS), and Nantes Université. The authors acknowledge the Cytocell -Flow Cytometry and FACS core facilty (SFR Bonamy, BioCore, Inserm UMS 016, CNRS UAR 3556, Nantes, France) for its technical expertise and help, member of the Scientific Interest Group (GIS) Biogenouest and the Labex IGO program supported by the French National Research Agency (n°ANR-11-LABX-0016-01).

## Appendix A. Supplementary data

## Data availability

Data will be made available on request.

