## supplementary file for "Ultrafast tyrosine-based cell membrane modification *via* diazonium salts: a new frontier for biomedical applications"

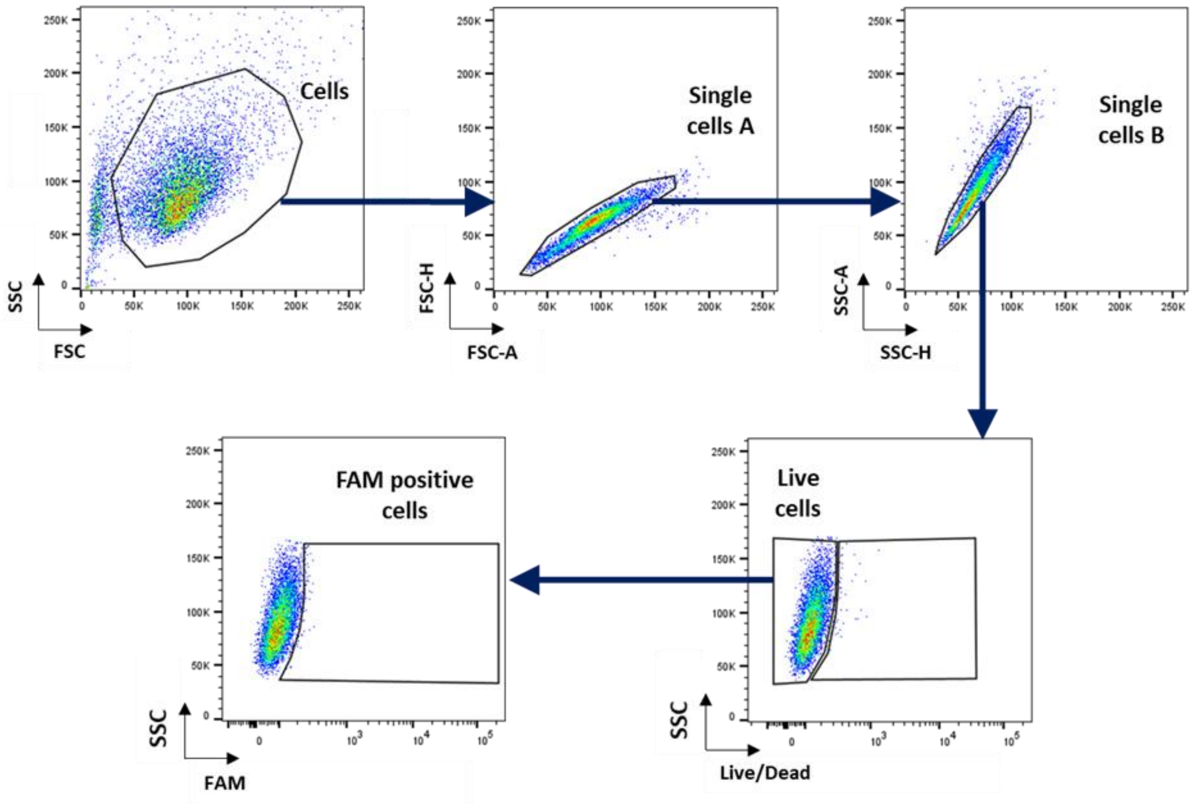


**Figure S1. General gating strategy for the different modified cell types.** Representative flow cytometry plots showing gating strategy after selection of the cell population. Single cells were identified by plotting forward scatter-area (FSC-A) against forward scatter-height (FSC-H), then by plotting side scatter area (SSC-A) against side scatter-height (SSC-H). Following dead cells staining using the LIVE/DEAD Viability/Cytotoxicity staining kit, FAM detection was performed on live single cells.

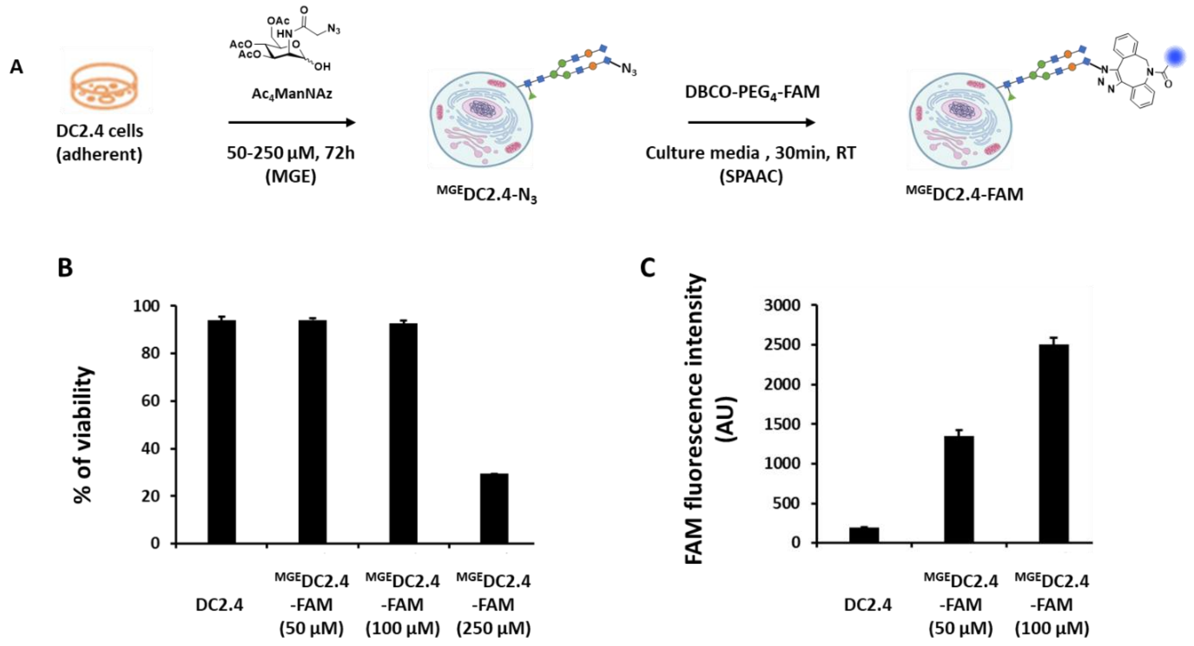


**Figure S2: DC2.4 cell membrane modification with MGE. A**) MGE protocol: DC2.4 cells in P24 were incubated with $Ac_4ManNaz$, followed by a 30 min SPAAC reaction with DBCO-PEG$_4$-FAM. **B)** Viability % of unmodified DC2.4 cells, DC2.4 cells incubated with $Ac_4ManNaz$ at 50, 100 or 250 µM for 72h, followed by a 30 min SPAAC reaction with DBCO-PEG$_4$-FAM (100 µM) ($^{MGE}$DC2.4-FAM (100µM), $^{MGE}$DC2.4-FAM (100µM) and $^{MGE}$DC2.4-FAM (250µM)) as evaluated by flow cytometry after incubation with LIVE/DEAD® assay kit ($n = 3$). **C)** FAM fluorescence intensity in arbitrary unit (AU) measured by flow cytometry for the first three samples ($n = 3$).

For DC2.4 cells were plated in a 24-well plate, and varying concentrations of $Ac_4ManNAz$ (50 to 250 µM) were added to the culture media. After 72 hours of incubation, the media was carefully aspirated, and the SPAAC reaction was performed under the same conditions as previously described. Our results demonstrated that $Ac_4ManNAz$ concentrations exceeding 100 µM ($^{MGE}$DC2.4-FAM) induced significant toxicity in DC2.4 cells, reducing cell viability to 30% at 250 µM.

Furthermore, we confirmed the efficacy of the MGE approach for chemically modifying the DC2.4 cell membrane. Indeed, an increase in fluorescence intensity was observed following incubation with DBCO-PEG$_4$-FAM, rising from 250 in untreated cells or cells incubated only with the fluorescent probe to 1500 at 50 µM and 2500 at 100 µM $Ac_4ManNAz$.

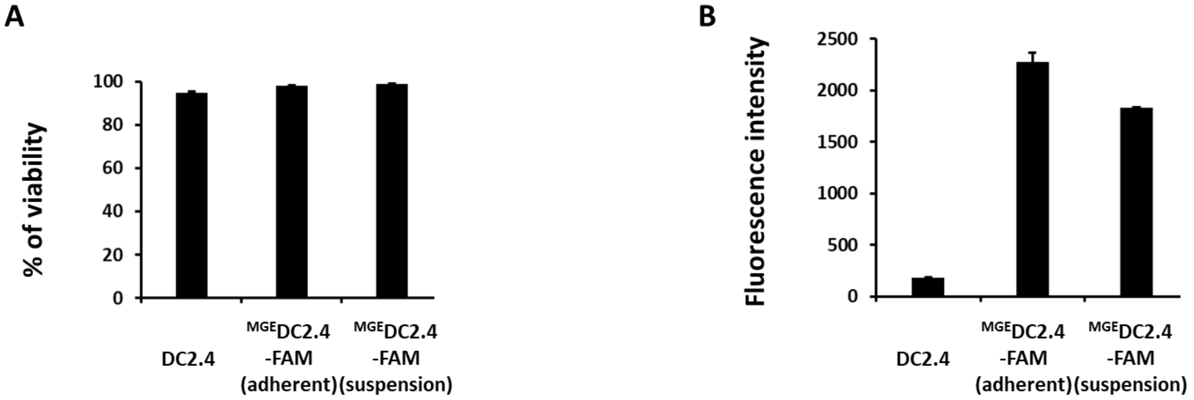


**Figure S3: Comparison of MGE with DC2.4 cells adherent and maintained in suspension. A**) Viability % of unmodified DC2.4 cells, DC2.4 cells incubated with $Ac_4ManNaz$ at 100 µM for 72h, followed by 30 min SPAAC with DBCO-PEG$_4$-FAM (100 µM) ($^{MGE}$DC2.4-FAM) in P24 (adherent), same protocol in low binding tube $^{MGE}$DC2.4-FAM (suspension), as evaluated by flow cytometry after incubation with LIVE/DEAD® assay kit ($n = 3$). **C)** Fluorescence intensity measured by flow cytometry for the four same samples ($n = 3$).

**A**

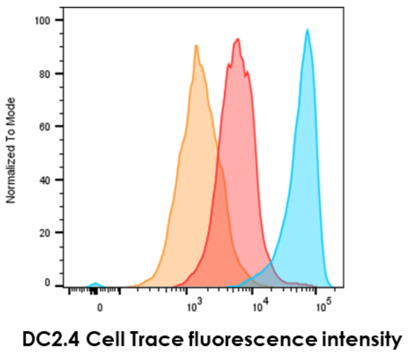


| Time | Fluorescence intensity |
|---|---|
| 0h | 61814 |
| 24h | 7060 |
| 48h | 2024 |

**B**

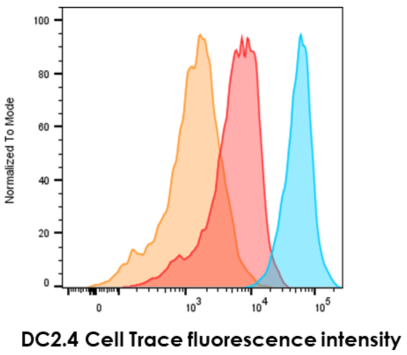


| Time | Fluorescence intensity |
|---|---|
| 0h | 59594 |
| 24h | 6960 |
| 48h | 1847 |

**Figure S4: DC 2.4 cells proliferation. A**) DC2.4 cells were incubated with CellTrace™ Far Red and fluorescence intensity measured by flow cytometry after 0h, 24h and 48h of cell culture ($n = 3$). **B**) DC2.4-$N_3$ cells (modified with **1** (100 µM)) were incubated with CellTrace™ Far Red and fluorescence intensity measured by flow cytometry after 0h, 24h and 48h of cell culture ($n = 3$).

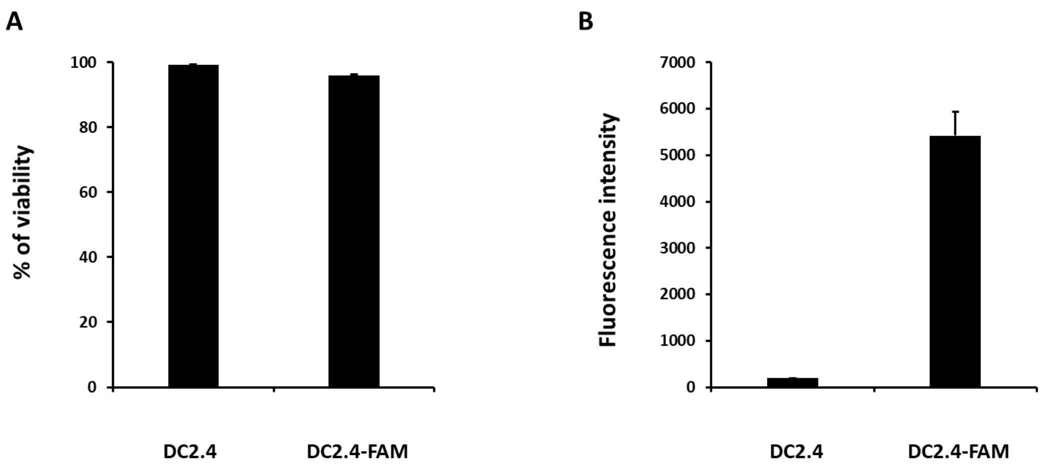


**Figure S5: Effect thawing on DC2.4-$N_3$ cells. A)** Viability % of unmodified DC2.4 cells and DC2.4-FAM cells after (incubation of DC2.4-$N_3$ cells (modified with **1** (5', 100µM) then with DBCO-$PEG_4$-FAM (30', 100µM)) after thawing as evaluated by flow cytometry after incubation with LIVE/DEAD® assay kit (n=3). **B**) Fluorescence intensity measured by flow cytometry for the two same samples (n=3).

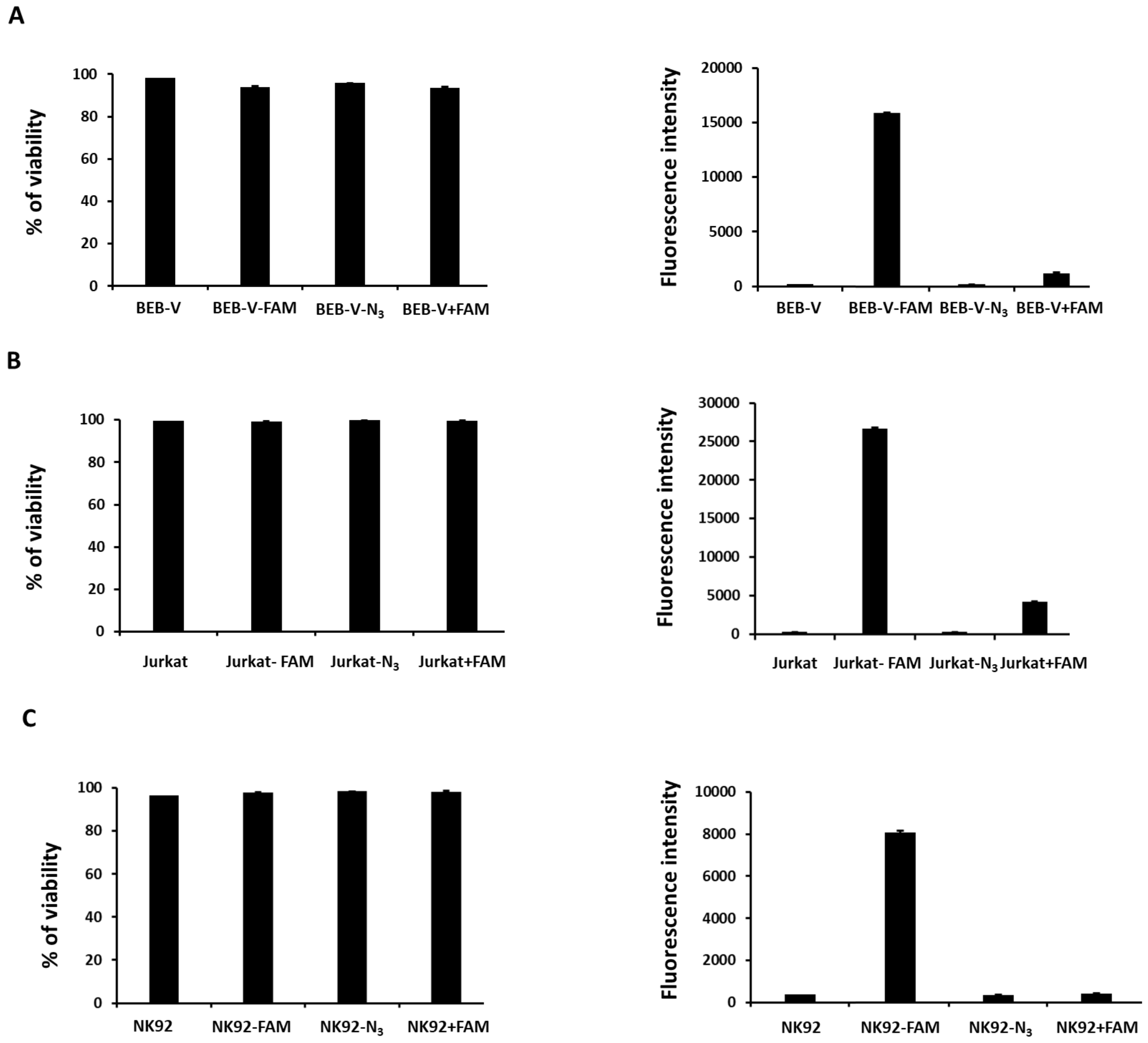


**Figure S6: Bioconjugation and SPAAC reaction to modify Y on protein of BEB-V, Jurkat and NK92 cell lines membrane. A**) Viability % and fluorescence intensity evaluated by flow cytometry, after incubation with LIVE/DEAD® assay kit, of unmodified B-EBV cells, BEB-V-FAM cells (modified with **1** (100 µM) and incubated with DBCO-PEG$_4$-FAM (100 µM)), BEB-V-N$_3$ cells (modified with **1** (100 µM)) and BEB-V + FAM cells (incubated only with DBCO-PEG$_4$-FAM (100 µM)) (n=3). **B**) Viability % and fluorescence intensity evaluated by flow cytometry, after incubation with LIVE/DEAD® assay kit, of unmodified Jurkat cells, Jurkat-FAM cells (modified with **1** (100 µM) and incubated with DBCO-PEG$_4$-FAM (100 µM)), Jurkat-N$_3$ cells (modified with **1** (100 µM)) and Jurkat + FAM cells (incubated only with DBCO-PEG$_4$-FAM (100 µM)) (n=3). **C**) Viability % and fluorescence intensity evaluated by flow cytometry, after incubation with LIVE/DEAD® assay kit, of unmodified NK92 cells, NK92-FAM cells (modified with **1** (100 µM) and incubated with DBCO-PEG$_4$-FAM (100µM), NK92-N$_3$ cells (modified with **1** (100 µM)) and NK92 + FAM cells (incubated only with DBCO-PEG$_4$-FAM (100 µM)) (n=3).

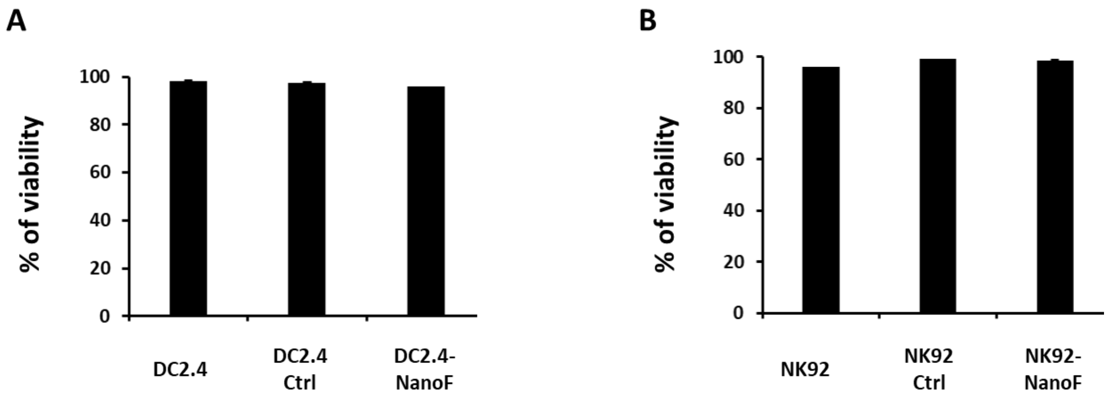


**Figure S7: Cell viability after Nanofitin coupling on Y of DC2.4 and NK92 protein cell membrane. A**) Viability % of non-modified DC2.4 cells, DC2.4 cells incubated with an Alexa-488 anti-histidine tag antibody (DC2.4 Ctrl), DC2.4-NanoF cells (modified with **1** (100 µM), incubated with DBCO-Nanofitin (50 µM), and incubated with an Alexa-488 anti-histidine tag antibody, as evaluated by flow cytometry after incubation with LIVE/DEAD® assay kit (n=3). **B**) Viability % of non-modified NK92 cells and the same others conditions as evaluated by flow cytometry after incubation with LIVE/DEAD® assay kit (n=3).

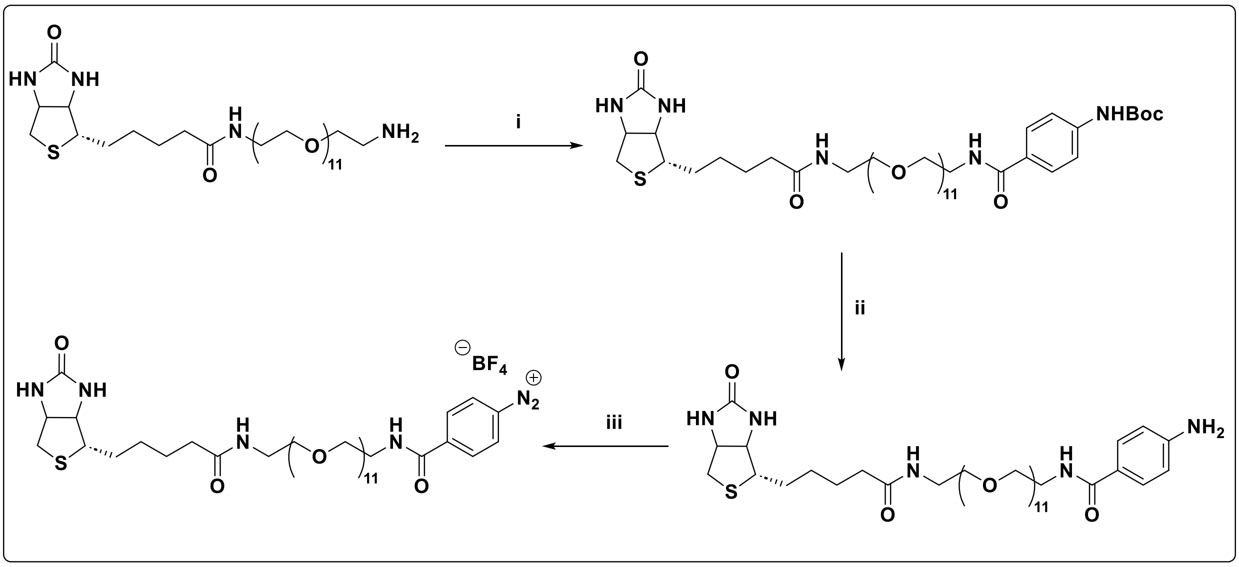


**Figure S8: Synthesis of biotin-$N_2^+$.** i) BocNHBzNHS, DIPEA, 1,4-dioxane, RT (86%), ii) DCM/TFA, 0°C (98%), iii) $HBF_4$, tBuONO, $H_2O$, RT (100%).

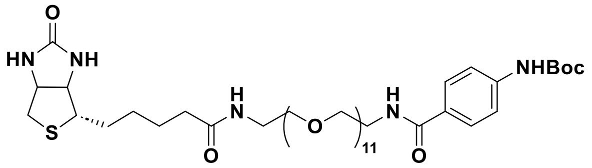


To a solution of biotin-$PEG_{11}NH_2$ (50 mg, 0.0649 mmol) in 1,4-dioxane (1.3 mL) were added BocHN-BzNHS (43 mg, 0.1298 mmol) and DIPEA (34 µL, 0.1947 mmol). After 12 h of stirring at 60°C, the solvent was evaporated under reduce pressure and the residue was purified by flash chromatography ($SiO_2$, DCM/MeOH: 95/5 → 90/10 → 80/20 as gradient eluents) to yield the amino-boc protected amine biotin derivative (55 mg, 0.055 mmol, 86 %) as a white solid.

$^1$H NMR ($CDCl_3$, 400 MHz): 1.40 (m, 2H), 1.49 (9H, s, NHBoc), 1.65 (m, 4H), 2.19 (t, 3H, *J* = 7.5 Hz, **$CH_2$**$CONHCH_2CH_2O$), 2.71 (d, 1H, *J* = 12.8 Hz), 2.86 (dd, 1H, *J* = 12.7 Hz, *J* = 4.8 Hz), 3.11 (m, 1H), 3.39 (m, 2H), 3.53 (t, 2H, *J* = 12.7 Hz, $CH_2CONH$**$CH_2$**$CH_2O$) 3.57-3.65 (m, 48H, PEG), 4.27 (dd, 1H, *J* = 7.7 Hz, *J* = 4.6 Hz), 4.49 (m, 1H), 5.67 (s, NH), 6.38 (s, NH), 6.80 (t, 1H, *J* = 5.4 Hz, NH), 7.05 (t, 1H, *J* = 5.4 Hz, NH), 7.33 (s, NH), 7.44 (d, 2H, *J* = 8.7 Hz, **Ph**NHBoc), 7.74 (d, 2H, *J* = 8.7 Hz, **Ph**NHBoc); $^{13}$C NMR ($CDCl_3$, 400 MHz): 25.6 ($CH_2$), 28.3 ($CH_2$), 28.4 (3 x $CH_3$, NHBoc), 29.8 ($CH_2$), 35.9 ($CH_2$), 39.2 ($CH_2$), 39.8($CH_2$), 40.6 ($CH_2$), 55.6 (CH), 60.3 (CH), 61.9 (CH), 69.9-70.6 ($CH_2$, PEG), 80.8 (C, NHBoc), 117.8 (2 x

CH, Ph), 128.3 (2 x CH, Ph), 128.6 (C, Ph), 141.8 (C, Ph), 152.7 (C, CO), 164.1 (C, CO), 167.1 (C, CO), 173.5 (C, CO); HRMS (ESI) for $C_{46}H_{80}N_5O_{16}S$ $[M+H]^+$, calcd 990.5321 found 990.5275.

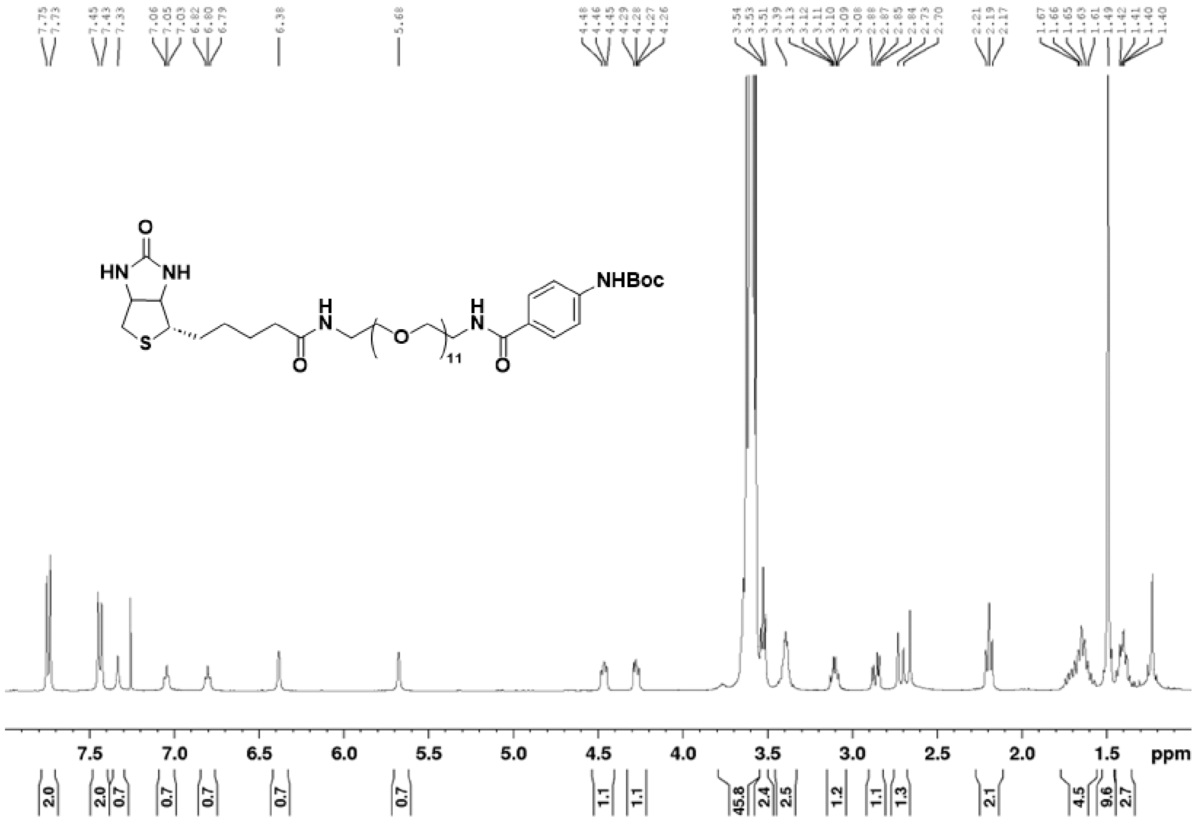


**Figure S9: RMN [1]H of the amino-boc protected biotin derivative**

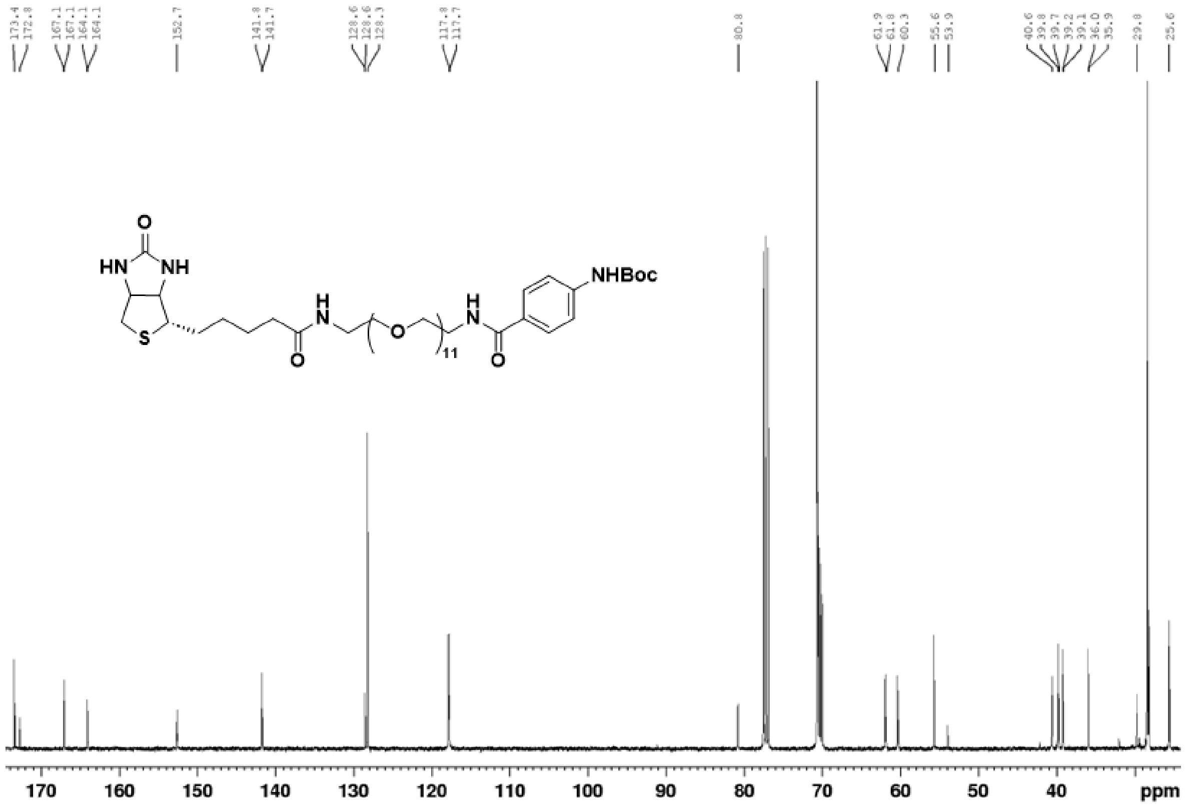


**Figure S10: RMN $^{13}$C of the amino-boc protected biotin derivative**

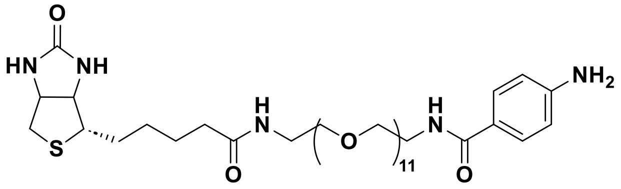


To a solution of the amino-boc protected biotin derivative (51 mg, 0.0515 mmol) in DCM (1 mL) was added, at 0°C, TFA (1 mL). After 1 h of stirring at 0°C, the solvents were evaporated under reduce pressure with co-evaporation with toluene (3 X) and the residue was lyophilized to yield the crude of the aniline biotin derivative (48 mg, 0.051 mmol, 98%) as a colorless oil. The crude of the reaction was used in the next step without further purification.

$^1$H NMR ($CD_3OD$): 1.42-1.79 (m, 6H), 2.23 (t, 3H, *J* = 7.5 Hz, $\mathbf{CH_2}CONHCH_2CH_2O$), 2.70 (d, 1H, *J* = 12.7 Hz), 2.92 (dd, 1H, *J* = 12.7 Hz, *J* = 5.0 Hz), 3.20 (m, 1H), 3.36 (t, 2H, *J* = 5.5 Hz, $CH_2CONH\mathbf{CH_2}CH_2O$), 3.52-3.76 (m, 53H), 4.29 (dd, 1H, *J* = 7.9 Hz, *J* = 4.5 Hz), 4.49 (ddd, 1H, *J* = 7.9 Hz, *J* = 5.0 Hz, *J* = 0.7 Hz), 6.79 (d, 2H, *J*= 8.9 Hz, **Ph**$NH_2$), 7.67 (d, 2H, *J*=

8.9 Hz, **Ph**$NH_2$); $^{13}C$ NMR ($CDCl_3$, 400 MHz): 25.5 ($CH_2$), 28.1 ($CH_2$), 28.4 ($CH_2$), 35.3 ($CH_2$), 38.9 ($CH_2$), 39.4 ($CH_2$), 39.7 ($CH_2$), 55.6 (CH), 60.2 (CH), 61.9 (CH), 69.2-70.1 ($CH_2$, PEG), 114.7 (2 x CH, Ph), 123.5 (C, Ph), 128.7 (2 x CH, Ph), 149.4 (C, Ph), 164.7 (C, CO), 168.8 (C, CO), 174.7 (C, CO); HRMS (ESI) for $C_{41}H_{72}N_5O_{14}S$ $[M+H]^+$, calcd 890.4796 found 890.4781.

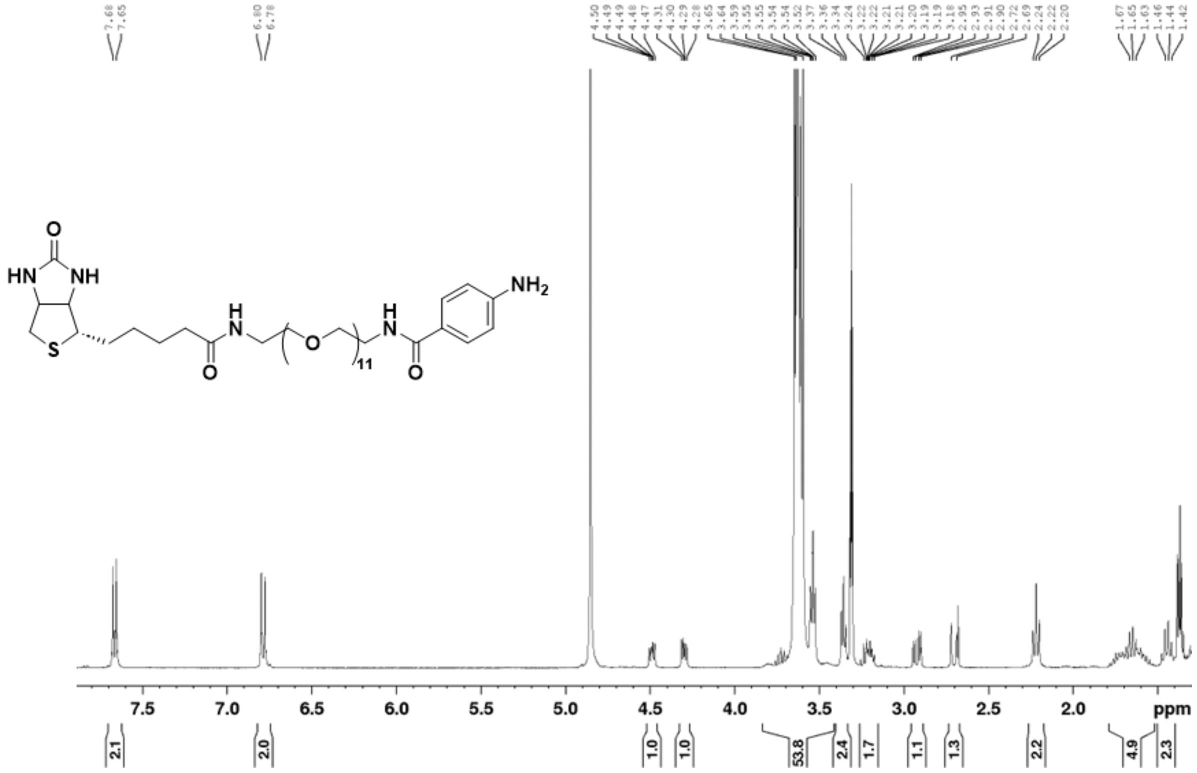


**Figure S11: RMN $^{1}H$ of the aniline biotin derivative**

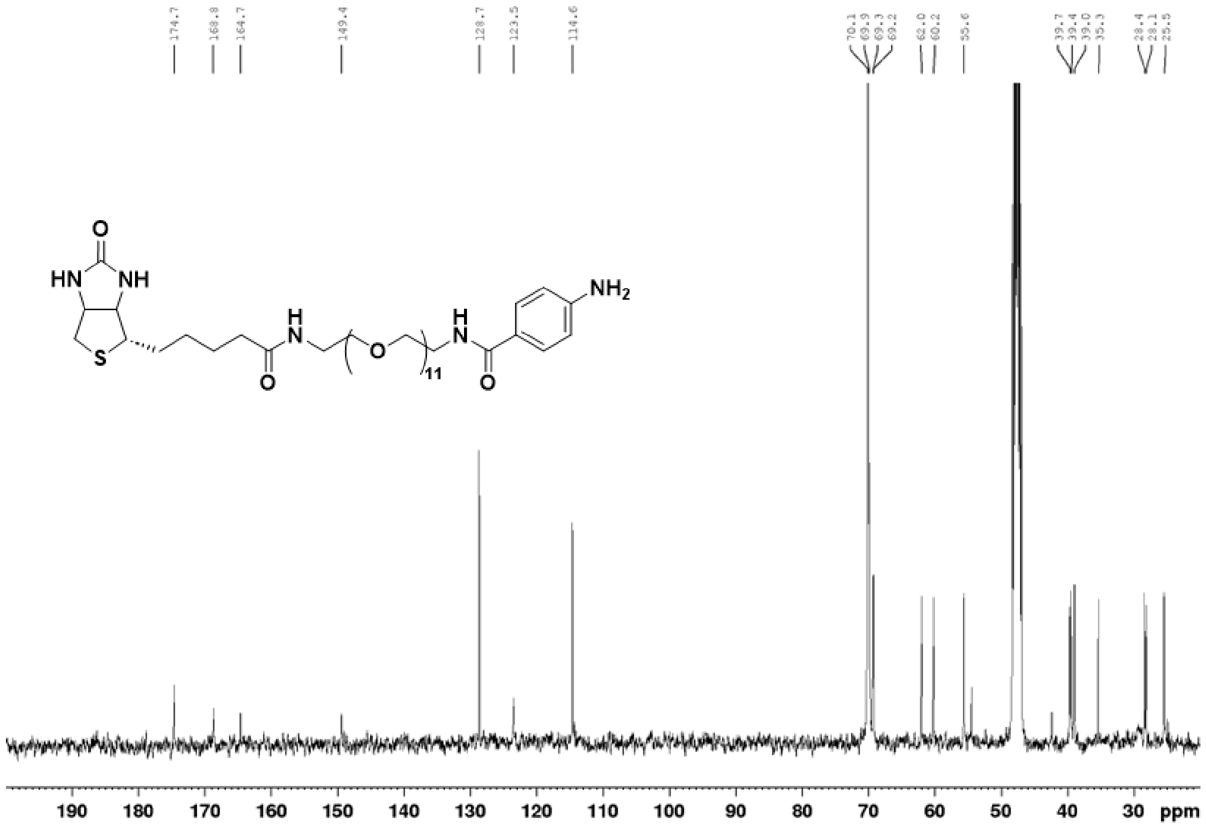


**Figure S12: RMN $^{13}C$ of the aniline biotin derivative**

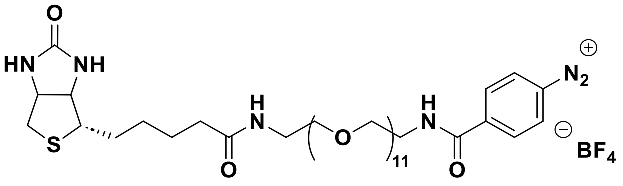

A solution of the aniline biotin derivative (51 mg, 0.0515 mmol) in $D_2O$ (0.8 mL) was placed in an NMR tube. Firstly, a solution of 50% $HBF_4$ in $H_2O$ (0.47 µL, 1 eq.) was added and the mixture was checked by 1H-NMR to monitor the aniline protonation. Next, tBuONO (0.31 µL, 1 eq.) was added and a second 1H-NMR was done to monitor the diazo compound formation.

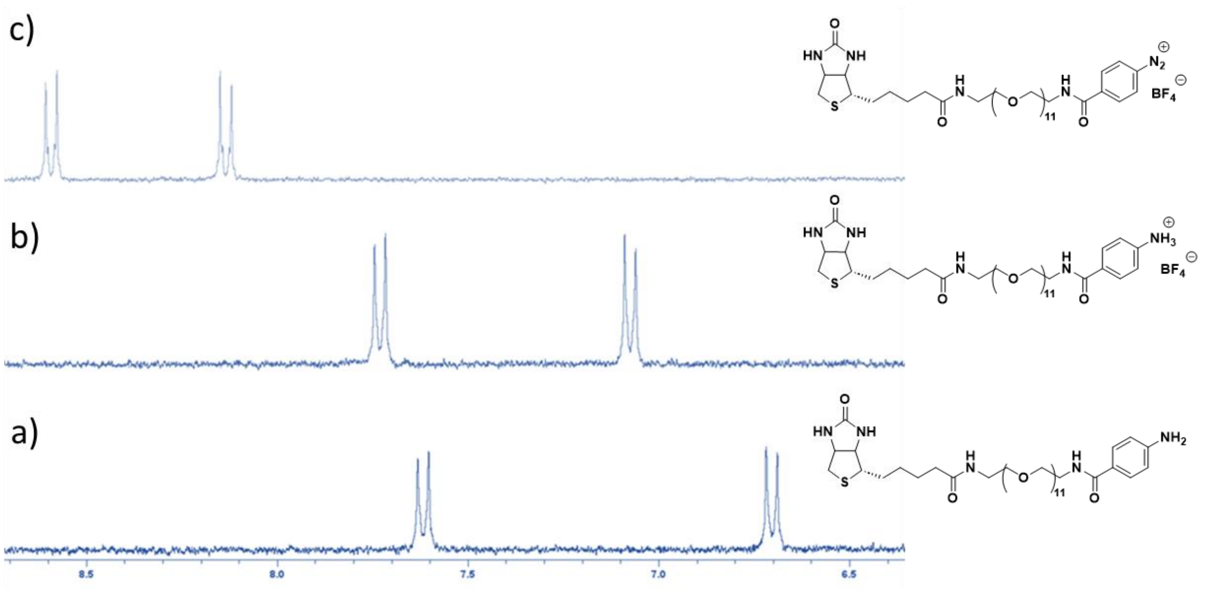


**Figure S13: ¹H-NMR monitoring formation of diazonium salt 2.** Monitoring of aromatic proton during diazo formation in $D_2O$: a) aniline, b) aniline after addition of 1 eq. of $HBF_4$, c) aniline after addition of 1 eq. of tBuONO.

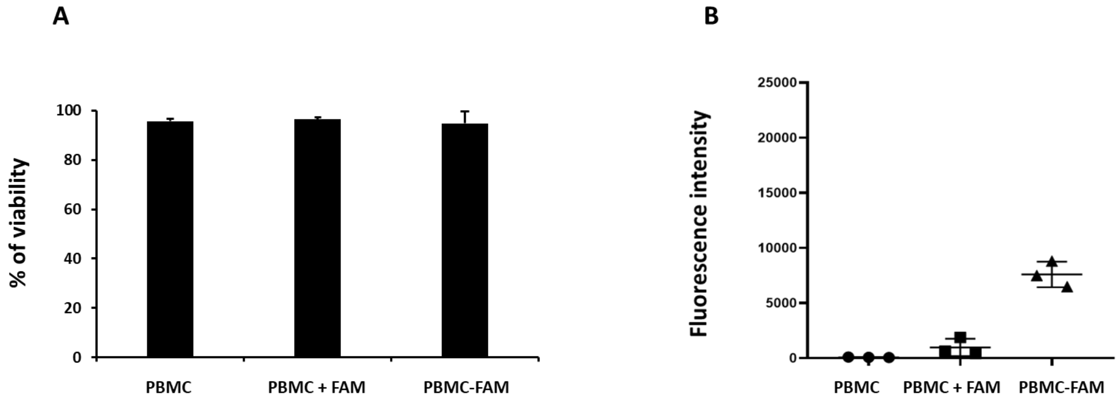


**Figure S14: Cell viability after DBCO-PEG$_4$-FAM coupling on Y of PBMCs protein cell membrane.** A) Viability % of unmodified PBMCs, PBMC-FAM cells (modified with **1** (100 µM) and incubated with DBCO-PEG$_4$-FAM (100 µM)) and PBMCs + FAM, cells (incubated only with DBCO-PEG$_4$-FAM (100 µM)), as evaluated by flow cytometry after incubation with LIVE/DEAD® assay kit. **B**) Fluorescence intensity of the same three samples as evaluated by flow cytometry of unmodified ($n = 3$).

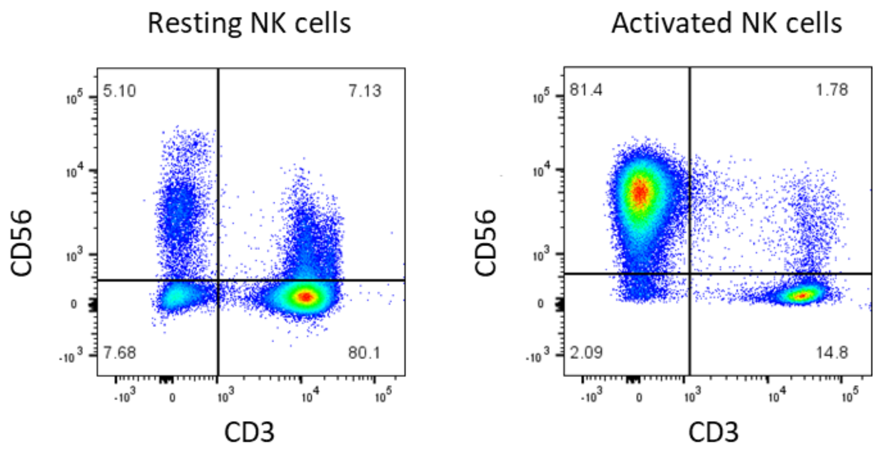


**Figure S15: Density plot of resting and activated NK cells.** Representative flow cytometry plots showing resting and activated NK cells identified as $CD3^{-}$ $CD56^{+}$ cells by flow cytometry using CD3 and CD56 specific mAbs.

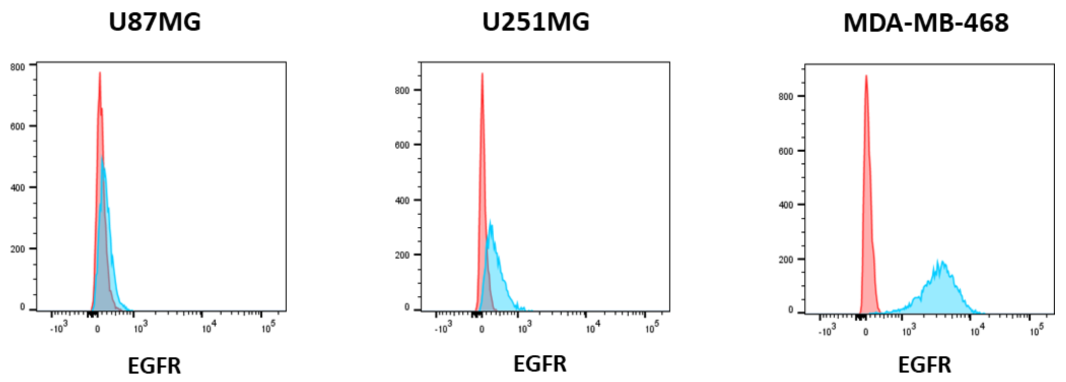


**Figure S16: EGFR expression on U87MG, U251MG and MDA-MB-468 cell lines.** EGFR expression on U87MG, U251MG and MDA-MB-468 cell lines was documented by flow cytometry using PE-EGFR-specific mAb.

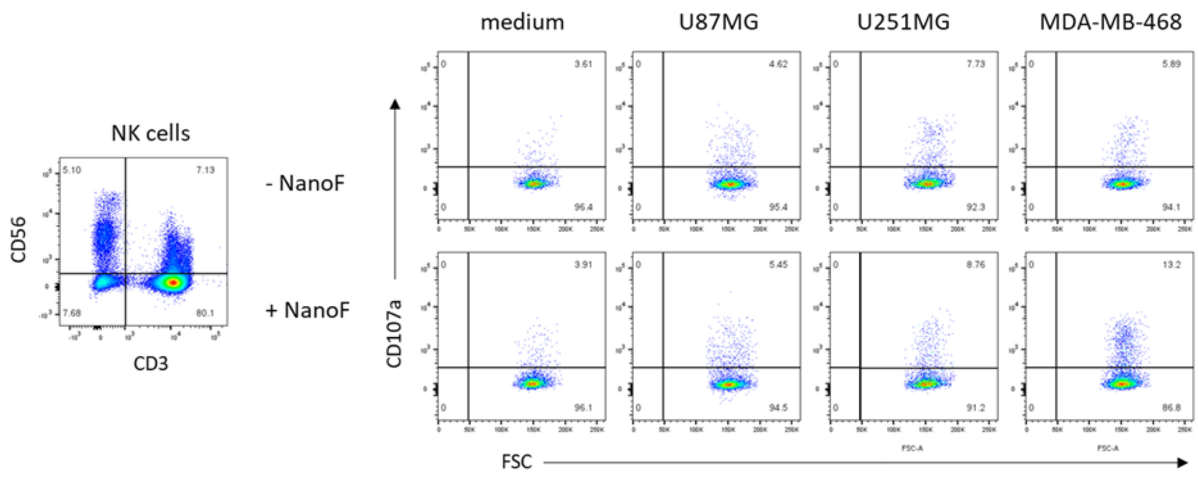


**Figure S17: Gating strategy of the degranulation assay performed on resting and activated NK cells.** Representative flow cytometry plots showing gating strategy and CD107a expression in resting NK cells and NK-NanoF cells after incubation for 5h with medium, U87M cell line, MDA-MB-468 cell line or U251MG cell line. Membrane staining was performed with CD3 and CD56 specific mAbs.
